# Man against machine: Do fungal fruitbodies and eDNA give similar biodiversity assessments across broad environmental gradients?

**DOI:** 10.1101/493312

**Authors:** Tobias Guldberg Frøslev, Rasmus Kjøller, Hans Henrik Bruun, Rasmus Ejrnæs, Anders Johannes Hansen, Thomas Læssøe, Jacob Heilmann-Clausen

## Abstract

The majority of Earths biodiversity is unknown. This is particularly true for the vast part of soil biodiversity, which rarely can be observed directly. Metabarcoding of DNA extracted from the environment (eDNA) has become state-of-the-art in assessing soil biodiversity. Also for fungal community profiling eDNA is seen as an attractive alternative to classical surveying based on fruitbodies. However, it is unknown whether eDNA-metabarcoding provides a representative sample of fungal diversity and census of threatened species. Therefore conservation planning and assessment are still based on fruitbody inventories. Based on a dataset of unprecedented width and depth, representing both soil eDNA-metabarcoding and expert inventorying of fungal fruitbodies, we document for the first time the validity of eDNA as practical inventory method and measure of conservation value for fungi. Fruitbody data identified fewer species in total and per site, and had larger variance in site richness. Focusing on macrofungi – the class Agaricomycetes, and in turn the order Agaricales – metrics of total richness and compositional similarity converged between the methods. eDNA was suboptimal for recording the non-soil dwelling fungi. β-diversity was similar between methods, but more variation in community composition could be explained by environmental predictors in eDNA data. The fruitbody survey was slightly better in finding red-listed species. We find a better correspondence between biodiversity indices derived from fungal fruitbodies and DNA-based approaches than indicated in earlier studies. We argue that (historical) fungal community data based on fruitbody forays – with careful selection of taxonomic groups – may be interpreted together with modern DNA-based approaches.

## 1. Introduction

### 1.1 methods for inventorying fungi

For decades, inventory and identification of fungal fruitbodies were – together with isolation and culturing – the only way to assess fungal communities (Hueck, 1953; Lange, 1948; Kjøller and Struwe, 1980; Rayner and Todd, 1980; Tyler, 1985; Schmit and Lodge, 2005). Since the 1990s, these methods have been supplemented with DNA-based methods, e.g. sequencing root samples to identify mycorrhizal fungi (Gardes and Bruns, 1996; Helgason *et al*., 1998) or sequencing of cloned PCR products from soil/litter samples (Schadt *et al*., 2003; O’Brien *et al*., 2005; Taylor et al 2014) – methods that allow for a more targeted study of some compartments, but are difficult to apply to ecosystem-wide inventories of large sampling sites. More recently, massive parallel sequencing of environmental DNA (eDNA) – now known as eDNA-metabarcoding (Taberlet *et al*., 2012) – has gained ground in studies of fungal communities (e.g. Schmidt *et al*., 2013; Pellissier *et al*., 2014; Tedersoo *et al*., 2014; Barnes *et al*., 2016), and allows for such wide inventories. In this study we compare a thorough fruitbody inventory with eDNA-metabarcoding for ecosystem-wide inventorying of the fungal community. Fruitbody surveys and eDNA-based methods both have their strengths and limitations and may be seen as complementary, rather than competing approaches (Truong *et al*., 2017).

### 1.2 Fruitbody inventorying

Fruitbody surveys are low tech but laborious, requiring life-long expert taxonomic skills if thorough and reproducible data are to be achieved (Newton *et al*., 2003). However, many fungi do not produce fruitbodies and are systematically omitted. Other taxa are likely to be under-sampled, as they are rarely fruiting, or produce very small, inconspicuous, short-lived or below-ground fruitbodies (Taylor and Finlay, 2003; Lõhmus 2009; van der Linde *et al*., 2012). Fruitbody formation and duration are highly influenced by (local variations in) season and weather conditions, which may hamper comparisons of sites, unless sampling is repeated over several years (Newton *et al*., 2003; O’Dell *et al*., 2004).

### 1.3 eDNA-metabarcoding

eDNA-metabarcoding is low tech when it comes to field sampling, but require high tech lab facilities and advanced post sequencing bioinformatics. It provides a broader taxonomic sample of the fungal community of not only the sexually reproducing, fruitbody forming fungi. Sampling of soil eDNA is less dependent on seasonality and climatic variation. Also, the majority of fungal biodiversity has yet to be described (Hibbett *et al*., 2011) and a large proportion of available barcode references lack proper annotation (Hibbett *et al*., 2011; Hibbett *et al*., 2016; Nilsson *et al*., 2016; Yahr *et al*., 2016). This limits ecological interpretation of detected community differences in relation to guild structure, trait space or taxonomic composition. Furthermore, when sampling for eDNA only a tiny fraction of a particular site surface area can be sampled, even with an intensive design. Hence, the sample representativity depends on the heterogeneity of species distributions within habitats and the size of mycelia (Lilleskov *et al*., 2004) – factors not easily assessed. This is a potential caveat, especially for detection of rare species – e.g. red-listed taxa – important for nature conservation (van der Linde *et al*., 2012). Although eDNA-metabarcoding has been shown to successfully identify red-listed species (Geml *et al*., 2014; van der Linde *et al*., 2012), these may be more easily detected as fruitbodies, which may be targeted by trained experts over large study areas in relatively short time, particularly for species with long-lived fruitbodies (e.g. perennial polypores; Runnel *et al*., 2015).

### 1.4 Fruitbodies versus eDNA-metabarcoding

Several studies detect a limited overlap in communities between fruitbody surveys and DNA-based approaches on a habitat scale (Gardes and Bruns, 1996; Dahlberg *et al*., 1997; Jonsson *et al*., 1999; Porter *et al*., 2008; Geml *et al*., 2009; Fischer *et al*., 2012; Baptista *et al*., 2015). However, it remains an open question whether key community metrics nonetheless correlate along environmental gradients, so that results from either method can be used as proxy for the other. In the context of nature conservation and monitoring, it would be attractive if eDNA-metabarcoding can be proven to detect target species (e.g. red-listed species) – for which e.g. historical data of decline is known, or where monitoring programs are already running – independent of optimal fruiting seasons and availability of taxonomic expertise. Finally, it would be valuable if historical data based on fruitbody surveys hold valid information on fungal communities, and may be compared or combined with modern eDNA-metabarcoding for inferring of temporal change.

### 1.5 Approach and expectations

In this study, we compare richness, community composition and community-environment relation in a large ecological space using two parallel data sets, a thorough fruitbody inventory and data obtained by eDNA-metabarcoding of soil. All data were gathered from the same 130 40×40 m sample plots in Denmark and taken over the same 2-3 year period.

Overall, we expected eDNA-metabarcoding to detect more species than fruitbody sampling. However, we expected the fruitbody survey to detect more red-listed species, due to the targeted survey across the whole study area and the fact that most red-listed species produce conspicuous fruitbodies. We expected eDNA-metabarcoding to provide stronger correlation with environmental gradients, due to the expected better coverage of taxonomic diversity. We expected comparability between the two approaches to be highest for community composition, and lowest for red-listed species detection, as previous studies have indicated stochastic variation in noisy data to affect richness estimates more than community composition (Abrego *et al*., 2016; Lekberg *et al*., 2014). Finally, we expected higher correspondence between fruitbody and eDNA-metabarcoding data, when the former were restricted to species recorded at soil level, and when both were restricted to Agaricomycetes and Agaricales.

## 2. Materials and methods

### 2.1 Study sites

130 sites of 40 × 40 m spread out across Denmark were studied. The study sites covered an ecospace spanning the major environmental gradients of terrestrial ecosystems, i.e. soil moisture, soil fertility and successional stage (Brunbjerg et al. 2017b). The 130 sites were selected by stratified random sampling to represent 24 environmental strata (habitat types). Six habitat types were cultivated: three types of fields (rotational, grass leys, set aside) and three types of forest plantations (beech, oak, spruce). The remaining 18 strata were natural habitat types, constituting all factorial combinations of: fertile and infertile; dry, moist and wet; open, tall herb/scrub and forest. We replicated these 24 strata in each of five geographical regions across Denmark. We further included a subset of 10 perceived biodiversity hotspots, two within each region. This study was part of the Danish biodiversity study, Biowide, and an elaborate description of design and data collection is available in Brunbjerg et al. (2017a).

### 2.2 Fruitbody survey

Each site was visited twice during the main fungal fruiting season in 2014 (August - early September and October - early November) and once during the main fruiting season in 2015 (late August - October), focussing on all groups of Basidiomycota and Ascomycota, but excluding non-stromatic Pyrenomycetes and Discomycetes with fruitbodies regularly smaller than 1 mm. Most woody debris was turned over to locate e.g. corticioid fungi, but no structured attempts to find hypogeous fungi were conducted, although a few were found by chance. In sites with tall and dense herbaceous vegetation, regular inspections were carried out in kneeling position. A site visit lasted approximately 1 hour, in very open monotonous sites sometimes less, e.g. in newly ploughed arable sites. All visits were led by expert field mycologist Thomas Læssøe typically accompanied by one helper. Numerous samples were taken back to a mobile lab for immediate microscopic investigation, and more interesting or critical material was dried as voucher material and in part deposited at the fungal herbarium (C) of the National History Museum of Denmark. Some specimens difficult to identify were forwarded to external experts.

### 2.3 Environmental variables

A complete inventory of vascular plants was done for each site. Ellenberg Indicator Values, EIV (Ellenberg *et al*., 1991) reflect plant species’ abiotic optimum and have often been used in vegetation studies to describe local conditions (Diekmann, 2003). Mean Ellenberg Indicator Values were calculated based on the plant lists for each site for the light conditions (EIV.L), soil nutrient status (EIV.N) and soil moisture (EIV.F). Ellenberg values together with measured variables (see supplementary methods) for precipitation, soil pH, soil organic matter content, soil carbon content, soil phosphorous and light were used in the models to explain community structure.

### 2.4 Sequence data

Soil was collected from all sites followed by DNA-extraction and sequencing with primers targeting fungi as described elsewhere (Brunbjerg *et al*., 2017a). For each site, 81 soil samples (each sample was approximately 5 cm diam. and 15 cm depth) were collected in a virtual grid with samples 4 m apart using a simple gardening tool. For each site, a large bulk soil sample was constructed by thorough mixing of 81 single soil samples, and subjected to DNA extraction with the MoBio PowerMax kit. The fungal ITS2 region was amplified using primers gITS7 (Ihrmark *et al*., 2012) and ITS4 (White *et al*., 1990). Libraries were MiSeq sequenced (Illumina Inc., San Diego, CA, USA), at the Danish National Sequencing Centre using two 250 bp PE runs. OTU tables (species-site table) were constructed, aiming for a definition of OTUs (Operational Taxonomic Units) that approximates species level delimitation. This was achieved by an initial processing with DADA2 (Callahan *et al*., 2016) to identify exact amplicon sequence variants including removal of chimeras, followed by ITS extraction with ITSx (Bengtsson-Palme *et al*., 2013) and subsequent clustering with VSEARCH (Rognes *et al*., 2016) at 98.5 % – the consensus clustering level used to delimit species hypotheses (SHs) in the UNITE database (Kõljalg *et al*., 2014), and subsequent post-clustering curation using LULU (Frøslev *et al*., 2017) to eliminate remaining redundant sequences. Taxonomic assignment of the OTUs was done using the 2017 UNITE general FASTA release (http://dx.doi.org/10.15156/BIO/587475). Sequence data is available from DataDryad, and files documenting the analyses from GitHub [[Data will be made available before publication. Until then it will be available by contacting the first author]].

For the more descriptive analyses, we used full fruitbody data. For some more direct comparisons, we restricted the fruitbody data to species collected at the soil surface for a more qualified comparison, as it was evident that only a small proportion of the non-soil fungi were registered by the soil-based eDNA-metabarcoding. Furthermore both datasets were filtered to obtain two increasingly taxonomically focussed subsets – Agaricomycetes and Agaricales. eDNA-metabarcoding and fruitbody data were then assessed for correspondence in a set of biodiversity metrics. Species composition was the focus of the study, and as biological abundance is difficult to assess with either method, presence/absence data was used for all analyses.

### 2.5 Overlap between methods

The frequency of each species/OTU across the 130 sites was assessed for the full datasets, and the proportion of species/OTUs recorded with both methods or only as fruitbody or OTU was assessed. As incomplete and insufficiently annotated DNA reference data exacerbate the discrepancies between fruitbody and metabarcoding data, some focussed analyses were performed on only the species recorded with both methods (‘coinciding species’).

### 2.6 Species richness and sampling effort

OTU richness was not greatly influenced by sequencing depth, Spearman rank r = 0.98 between OTU count based on rarefied data (10,000 reads per sample) and full data, and OTU richness measures were thus estimated from the full (not rarefied) data. Species accumulation was assessed for all datasets, and the variation of the recorded site richness per site was assessed by calculation of the relative standard deviation of richness for each method. Pearson correlation was used to test for correspondence between estimates of species richness and OTU richness across the 130 sites.

### 2.7 Red-listed species

As a measure of conservation value we used the count of red listed fungal species in the IUCN categories from near threatened to critically endangered on the official Danish red list (IUCN 2012, Wind & Pihl 2010). We assessed both the total number of red-listed species identified with either method, as well as the correspondence of site wise counts of red-listed species.

### 2.9 Community composition

Community dissimilarity was estimated with the Sørensen dissimilarity metric using the vegdist function in vegan. Five out of the 130 sites had less than 4 observed fruitbody species and were removed prior to analyses of community dissimilarity. Correlation between community dissimilarity measures based on different datasets were tested with the Mantel test (method = "pearson", 999 permutations) and Procrustes test (999 permutations) using the functions in vegan. Community turnover along gradients (assessed as dissimilarity) was tested for correlation with environmental distance using the *bioenv* function in vegan. Here Sørensen distance is used for community dissimilarity and Euclidean distance for environmental dissimilarity, and we allowed up to four explanatory variables to be selected.

## 3 Results

### 3.1 Overlap between methods

The fruitbody survey recorded fewer species than the eDNA-metabarcoding approach (Fig. 1a, Supplementary Fig. 1). The fruitbody survey included 8,793 observations (a species in a site), and recorded 1,751 species (1,358 Agaricomycetes of which 847 belonged to Agaricales). The eDNA-metabarcoding included total 30,668 observations (an OTU in a site), and recorded 8,110 OTUs (2,521 Agaricomycetes and 1,293 Agaricales). 1,288 (74 %) of the fruitbody species were recorded as fruitbodies only, while 463 (26 %) were found also as OTUs. 7517 (93 %) of the OTUs were found with eDNA-metabarcoding only, while 593 (7 %) were recorded also as fruitbodies (i.e. had species name annotations corresponding to the 463 species mentioned above). For these 463 coinciding species there was a tendency towards pairwise correspondence of species and OTU frequency (Fig. 1a and 1c). Four coinciding species were common as fruitbodies, but rare as OTUs. Three of these *(Mycena speirea, Clitopilus hobsonii, Mollisia cinerea)* are normally observed on woody or herbaceous substrates, and the last *(Galerina vittiformis)* is associated with bryophytes. The top ten most frequent coinciding OTUs were less frequent as fruitbodies – all common soil fungi, except *Ganoderma applanatum*, a wood decomposer not generally perceived as a soil fungus. Community dissimilarity estimates based on the 463 coinciding species resulted in corresponding composition estimates (mantel r-statistic 0.63, and 0.83 correlation in a symmetric procrustes rotation). Site OTU richness and fruitbody species richness was highly correlated when considering only the 463 coinciding species, r = 0.81 with a slope close to 1 (Fig. 1d).

**Figure 1.**
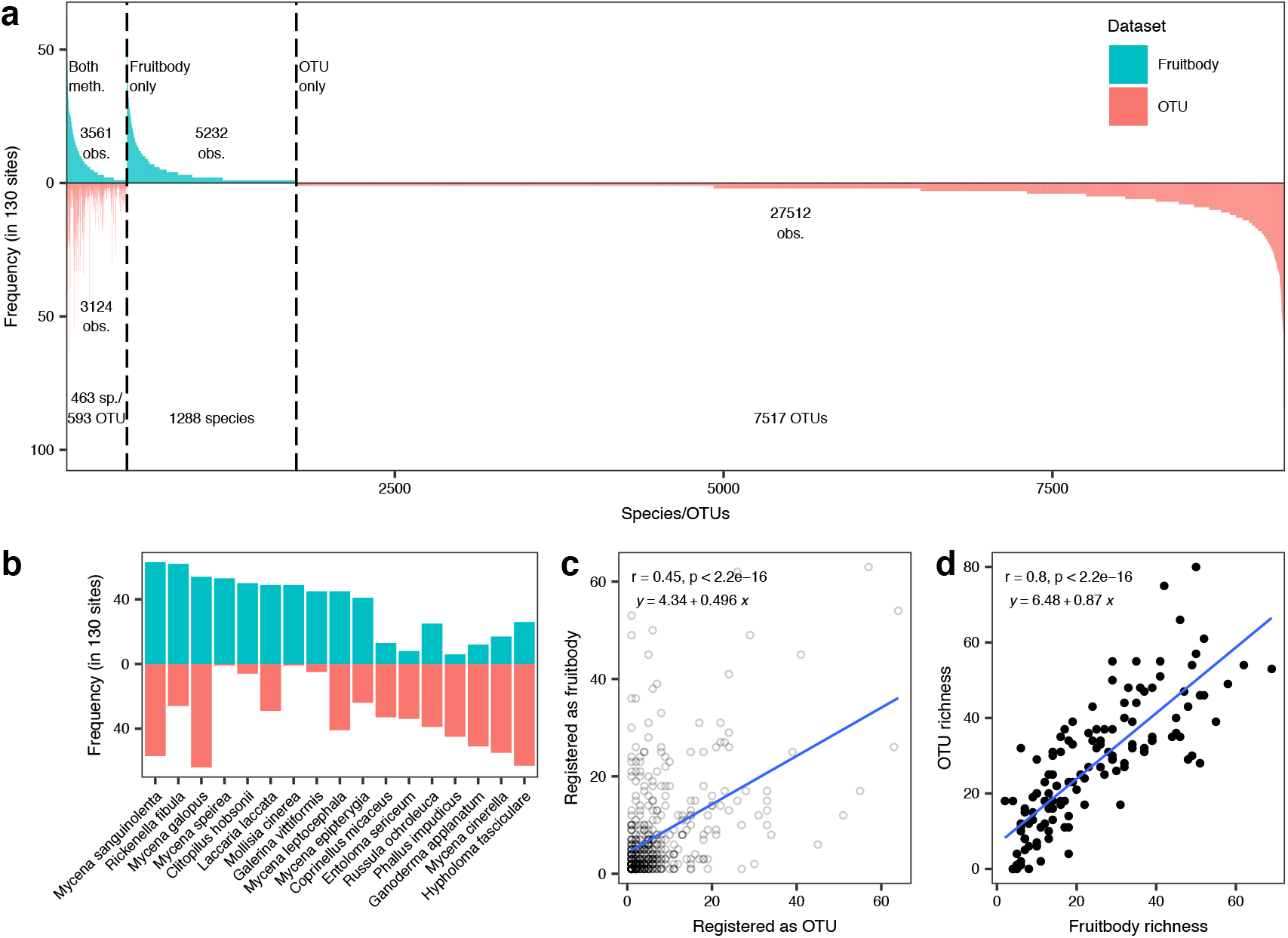
Frequency of species (and OTUs) among the 130 sampling sites. a) Frequency of species sorted by decreasing frequency, and grouped by species recorded with both methods or only as OTU or as fruitbody, y-axis indicates the number of sampling sites (of 130) in which a species was recorded, number of species and number of observations (a species/OTU in a site) are indicated for each group. b) Top 10 most frequent species recorded with either method. c) scatterplot of fruitbody-based frequency vs DNA frequency of the 463 species recorded by both methods. d) Species richness of the 130 sites as recorded with fruitbodies or OTUs for the 463 species recorded with both methods.

394 (37%) of the 1,067 soil fruitbody species were also registered as OTUs, whereas only 69 (10%) of the 684 non-soil fruitbody species were also registered as OTUs. Per site, an average of 6.3% of the soil fruitbody species were captured as OTUs, but only 0.18% of the non-soil fruitbody species. As it was evident that the soil eDNA captured little of the non-soil funga, we made comparison analyses of site richness and red-list recording both using the full fruitbody data but also on fruitbody data excluded the non-soil species.

### 3.2 Overall species richness and sampling effort

Although the exact logging of expenses was not part of the project, we estimate that the costs of the two approaches would be approximately equal, if repeated with the focussed aim of monitoring. The fruitbody survey included 3 × 10,000 km driving, and four months of salary (three months of collecting, one month of identification) – excluding the aid from volunteers in the fruitbody survey, whereas the eDNA-metabarcoding included 10,000 km driving, approximately 6,000 USD lab consumables, and 3 months salary (1 month collecting, 2 months lab work and bioinformatics). Species accumulation curves did not reach an asymptote for any of the datasets after sampling of the 130 sites (Fig. 2a). This was most pronounced for the full eDNA-metabarcoding dataset and least for the non-soil Agaricales fruitbody dataset. eDNA-metabarcoding became increasingly similar to fruitbody data with narrowed taxonomic focus. The variation in site species/OTU richness across the 130 sites was lowest for eDNA-metabarcoding, and highest for fruitbody data (markedly higher for non-soil fungi), but more similar with a narrowed taxonomic focus (Fig. 2b).

**Figure 2.**
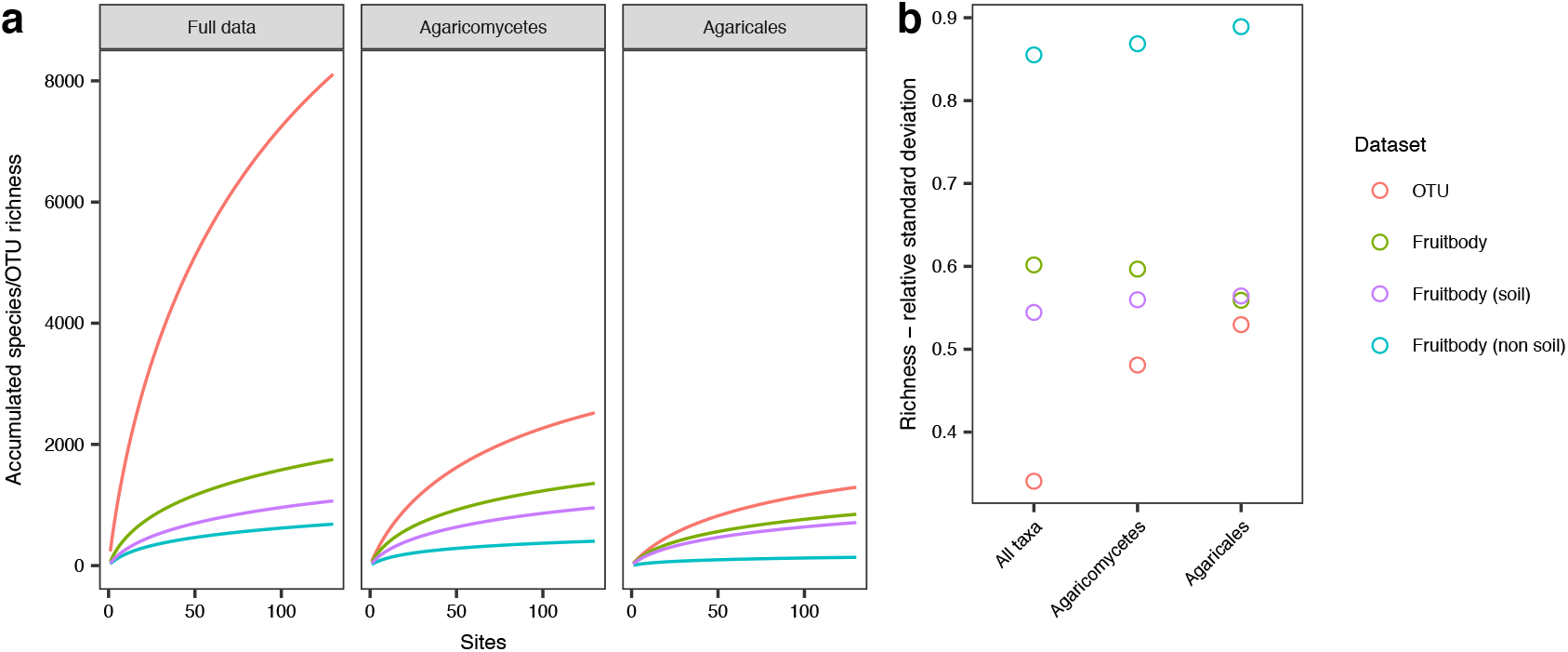
Sampling effort and richness variation. a) Cumulative species richness when sampling the 130 sites for the full data, Agaricomycetes and Agaricales. b) Relative standard deviation (RSD) of site species/OTU richness across the 130 sampling sites for the full data, Agaricomycetes and Agaricales.

### 3.3 Richness correlation between methods

Soil fruitbody species richness of the 130 study sites (Fig. 3) ranged from 0-115 (0-111 Agaricomycetes and 0-79 Agaricales), while OTU richness ranged from 66-476 (11-157 Agaricomycetes, 6-87 for Agaricales). Correlation between site species richness and OTU richness (Fig. 3) was moderate for the full datasets (r =0.43), but strong when restricted to Agaricomycetes (r = 0.64) and Agaricales (r = 0.58). Correlations became even stronger (r 0.74 – 0.81) when only considering the 463 ‘coinciding species’ – species recorded with both methods (Fig. 1d, Supplementary Fig. 1d and 1h). OTU richness based on only Agaricomycetes or Agaricales were strongly correlated with OTU richness based on the full data and (r = 0.74 and r = 0.8, respectively, Supplementary Fig. 2).

**Figure 3.**
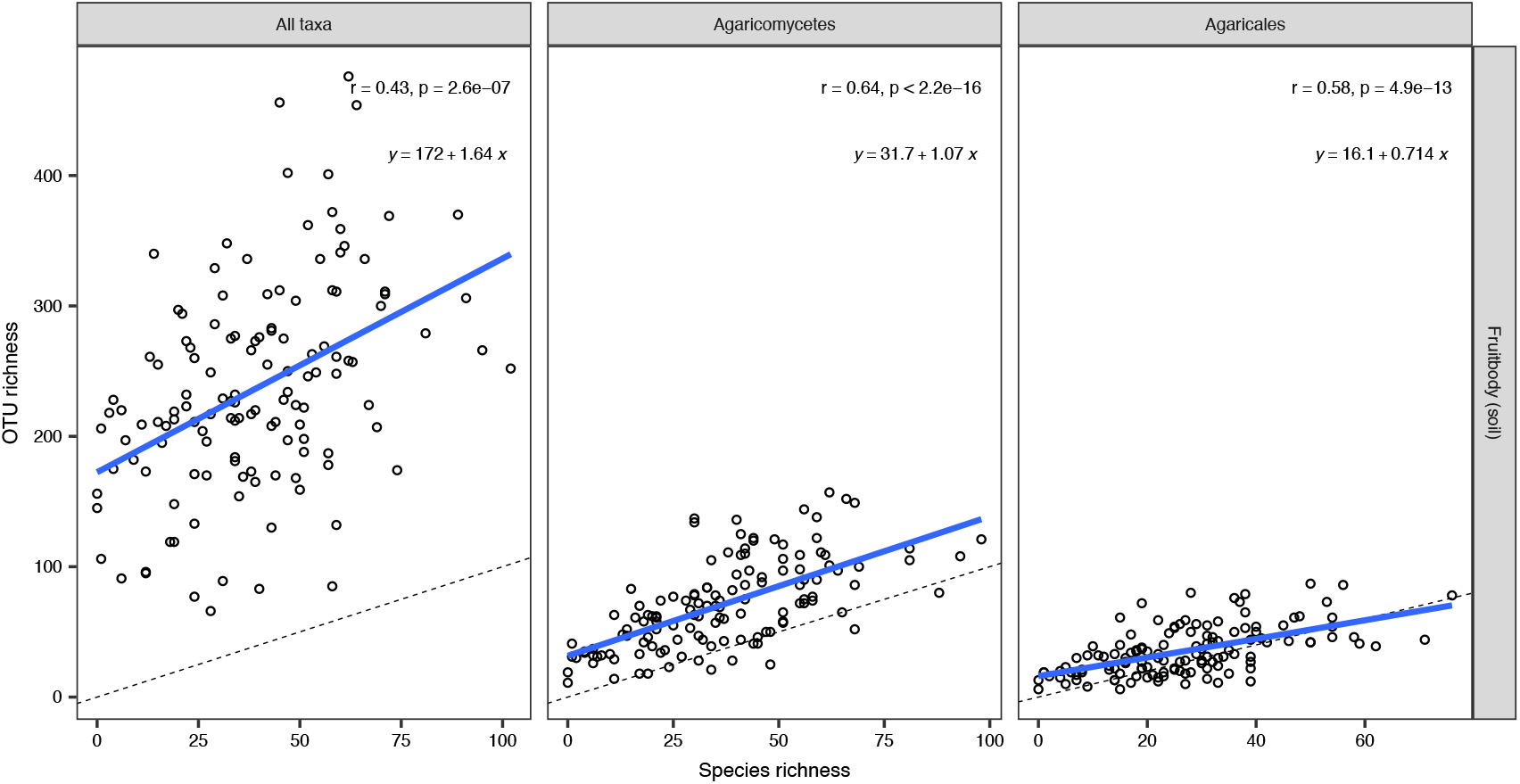
Correlation between site fruitbody species richness and OTU richness. Blue lines represent the linear regression of OTU richness against species richness, while the dotted line shows the identity line (x=y). Correlations are shown for the taxonomic subsets (all taxa, Agaricomycetes, Agaricales). Fruitbody data is restricted to species registered at the soil surface.

### 3.4 Overall taxonomic composition

Taxonomic composition of eDNA-metabarcoding and fruitbody data became increasingly similar when going from full data to Agaricomycetes and Agaricales (Fig. 4, Supplementary Fig. 3, Supplementary Tables 1-4). Fruitbody data was heavily skewed towards Basidiomycota (90 %), whereas the eDNA-metabarcoding was composed of 39 % Ascomycota, 40 % Basidiomycota, and 20 % species from other phyla (Fig. 4a, Supplementary table 2). However, the relative proportions and absolute frequencies of taxa progressively converged when focussing on Agaricomycetes (Fig. 4b, Supplementary Fig. 3c) and Agaricales (Fig. 4c, Supplementary Fig. 3d). The non-soil fruitbody data was less dominated by Agaricomycetes and Agaricales than the soil-fungi data. All phyla and classes (except Dacrymycetes and Atractiellomycetes) were represented by more species/OTUs in the eDNA-metabarcoding than in the fruitbody data (Fig. 4, Supplementary Fig. 3ab). A few Agaricomycetes orders (Polyporales, Hymenochaetales, Auriculariales, Gomphales) were represented by more species in the fruitbody data than in the eDNA-metabarcoding (Supplementary Fig. 3c). Almost all Agaricales genera were detected by both methods and with roughly similar species numbers.

**Figure 4.**
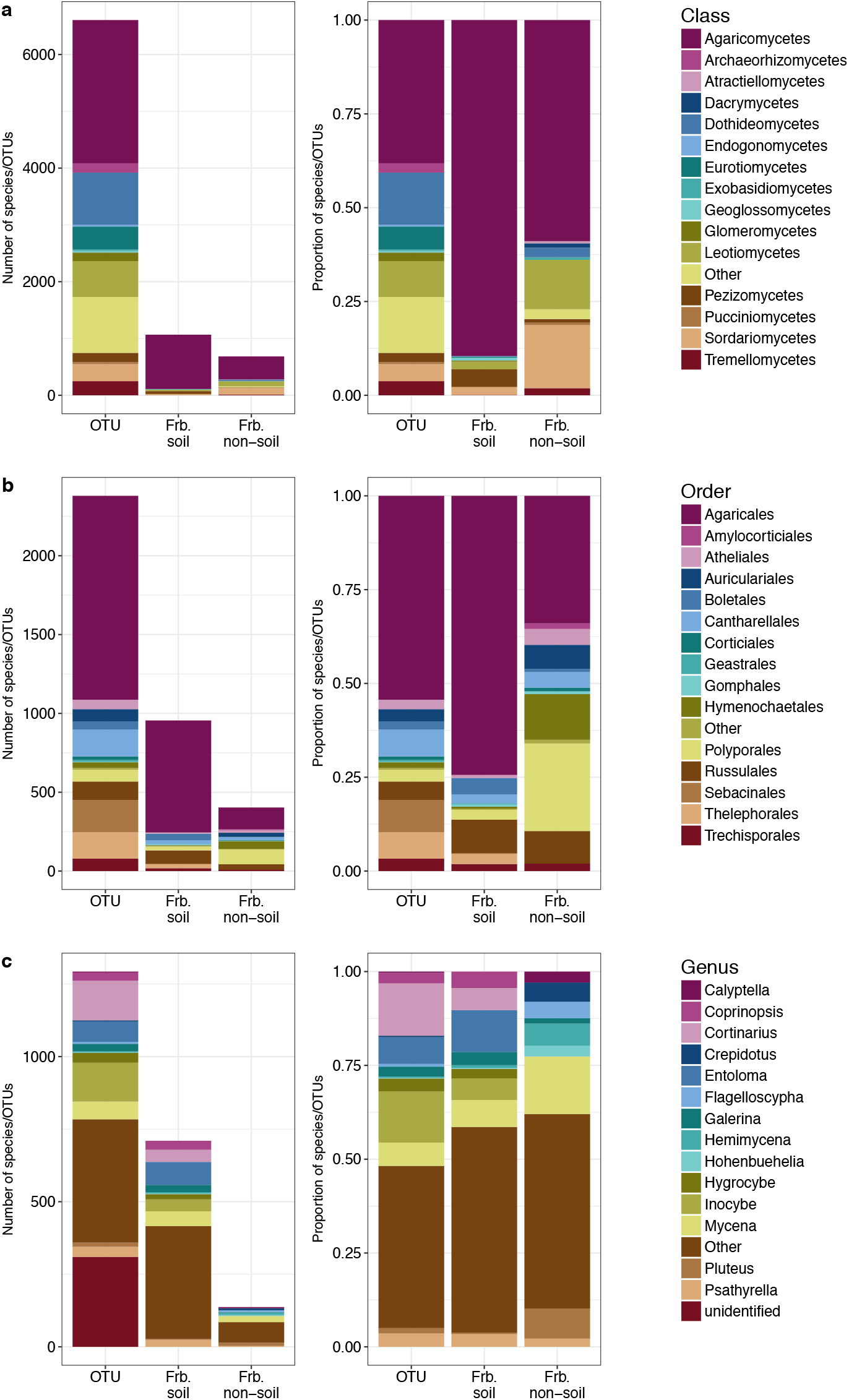
Taxonomic composition. Plots show the number of species (and OTUs) assigned to higher taxa. Composition is shown for OTUs, fruitbody (soil) and fruitbody (non-soil). Left plot in each panel shows the absolute richness (number of species) within different taxa, right plot shows the relative richness. a) Number of species in each class for full datasets. b) Number of species in each order in Agaricomycetes. c) Number of species in each genus of Agaricales. Most frequent taxa for each dataset is shown for all datasets, the rest are pooled in the category ‘Other’.

### 3.5 Red-listed species

The soil surface fruitbody survey recorded more red-listed species than the eDNA-metabarcoding (Fig. 5, Supplementary Table 5). 100 red-listed species were recorded as fruitbodies on the soil surface (144 including the non-soil fungi), whereas 63 red-listed species were found as OTUs. 26 red-listed species were recorded with both methods, 37 red-listed species was detected as OTUs only, and 74 as fruitbodies only. Only one red-listed species from the non-soil part of the fruitbody data (*Ganoderma pfeifferi*) was also detected as OTU. When restricting the comparison of red-listed species to species present in the molecular reference database the figures for soil fruitbodies (71species) was almost equal to the figures for OTUs (Fig. 5). When the 130 sites were grouped into categories with 0, 1-2 or 3 or more red-listed species recorded as fruitbodies or as OTUs, there was a good correspondence between the two methods (Supplementary Fig. 4).

**Figure 5.**
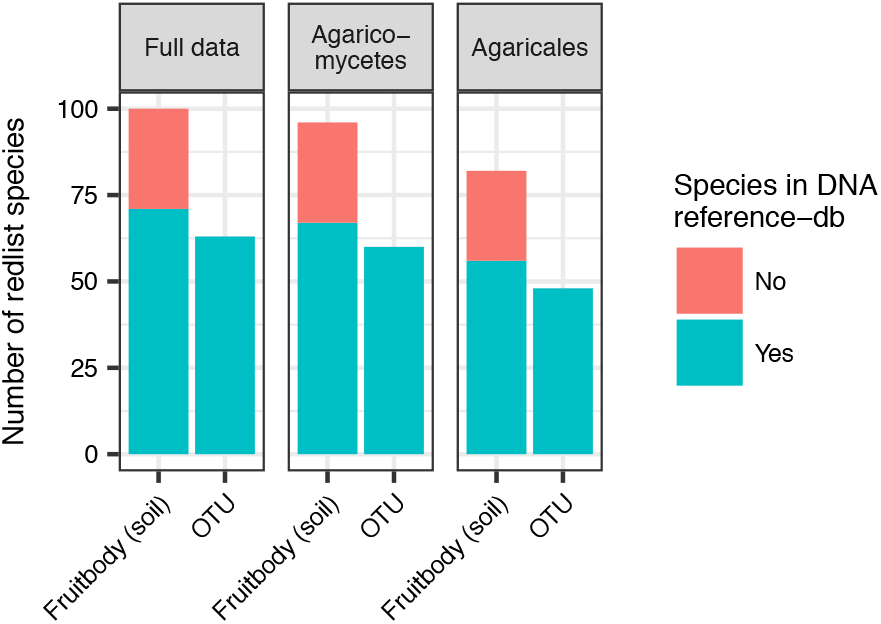
Number of red-listed species recorded. Total number of red-listed species found as fruitbodies (restricted to soil fungi), and OTUs. Taxa present in the DNA reference database (and thus possible to identify with both methods) are indicated in green, and taxa not present in the DNA reference database (and thus not possible to identify with DNA) are indicated in red.

### 3.6 Community – environment relation

Mantel tests showed very similar and strong correlations between community dissimilarity measures from fruitbody and eDNA-metabarcoding data, mantel r = 0.67. In this case no improvements were achieved by increased taxonomic overlap (Mantel-r = 0.68 and 0.62 for Agaricomycetes and Agaricales respectively), or when restricted to soil-fungi (Mantel-r = 0.68, 0.71 and 0.64 for full data, Agaricomycetes and Agaricales respectively). These correlations were corroborated by procrustes analyses with correlation coefficients of 0.87, 0.87 and 0.83, and 0.87, 0.87, 0.83 for soil-fungi for the same comparisons (all p-values < 0.01). Environmental variables explained more of the community dissimilarity for eDNA-metabarcoding data than for the survey data, and the amount of explained variation was largest for the taxonomically more inclusive datasets (Fig. 6) with the maximum explained variation for the full eDNA-metabarcoding dataset (0.66) and the lowest for the soil-fruitbody Agaricales (0.50). Adding a fourth explanatory variable did not increase the amount of explained variation for most datasets. Based on all subsets of the fruitbody data, the best three explanatory variables for community composition were mean Ellenberg soil nutrient status (EIV.N), mean Ellenberg light indicator value (EIV.L) and soil phosphorous, whereas mean Ellenberg soil nutrient status (EIV.N), mean Ellenberg soil moisture (EIV.F) and soil pH were the best for the DNA metabarcoding.

**Figure 6.**
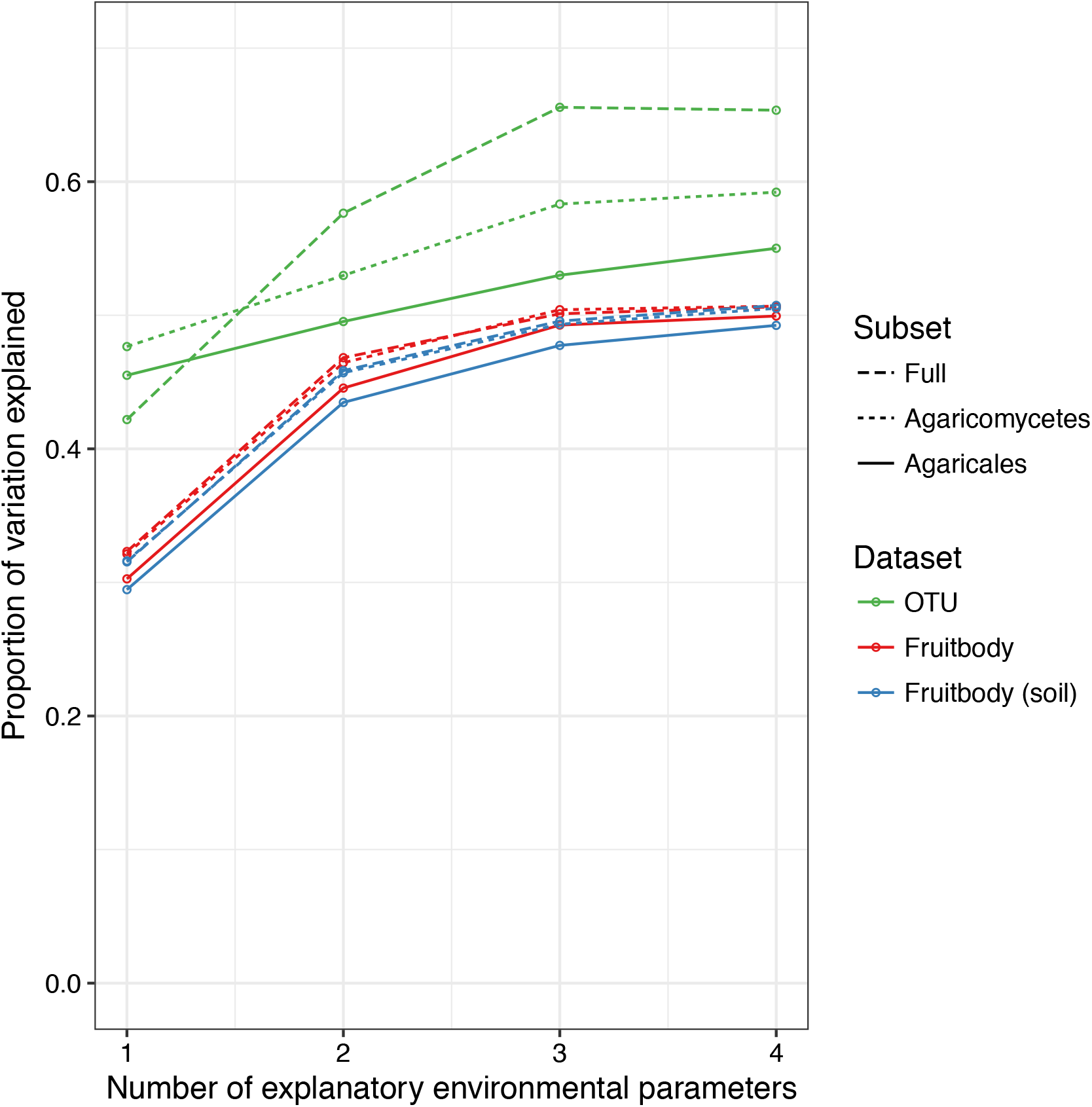
Proportion variation of community dissimilarity. X-axis shows the number of explanatory environmental variables selected by the model, and y-axis shows the total amount of explained variation. Colours indicate the dataset (OTU, fruitbody (all data) and fruitbody (soil species), line type indicates the three taxonomic subsets (All taxa, Agaricomycetes and Agaricales).

## 4. Discussion

More species (OTUs) were detected by eDNA-metabarcoding than by the classic fruitbody survey. This could mainly be attributed to the detection of groups, which always go undetected in a fruitbody survey, e.g. diverse groups of moulds and yeasts. The fruitbody survey data was strongly dominated by fruitbody-forming basidiomycetes. In general, there was a relatively poor correlation for richness measures and taxonomic composition between the two full datasets, but increased strength of correlation when narrowing the focus to Agaricomycetes and subsequently to just Agaricales. Similarly, excluding wood-inhabiting and other non-soil fungi improved the correspondence between the datasets, showing that these largely go undetected in soil-based eDNA sampling. The fruitbody survey identified more red-listed species, but the difference was less pronounced than anticipated, and results were almost similar when delimited to soil-dwelling fungi only.

### 4.1 Taxonomic composition similar for macrofungi

The taxonomic composition was remarkably similar between eDNA-metabarcoding and fruitbody data when focussing on the Agaricomycetes, and even more pronouncedly the Agaricales. Many of the major discrepancies align with expectations – i.e. taxonomically difficult groups like *Inocybe* (Larsson et al 2009, Ryberg et al. 2008) and *Cortinarius* (Frøslev et al. 2007) were markedly more species rich as eDNA OTUs than as well-delimitated species identified from fruitbodies. Approximately half of the Agaricales species recorded as fruitbodies were also found as OTUs and vice versa (Supplementary Fig. 1e). Considering the very similar proportions of Agaricales genera between the methods, it can be assumed that a large part of non-overlapping species can be explained by incomplete DNA reference data and different taxonomic concepts in handbooks for species identification of fruitbodies compared to sequence databases. An effort to expand and curate DNA reference databases is hence essential to improve future DNA-based ecological studies as already suggested by other researchers (Hibbett *et al*., 2011; Hibbett *et al*., 2016; Nilsson *et al*., 2016; Yahr *et al*., 2016).

### 4.2 Soil DNA captures soil-fungi

In this study, we extracted DNA from soil samples. Although DNA from non-soil fungi may be expected to be present in the soil, it has not earlier been tested to what extend soil DNA can be used to register fungi not having their active growing life-stages within the soil, such as wood decomposing fungi. Several non-soil fungi were detected in this study, but they were observed in much fewer sites than soil-fungi when comparing to the corresponding fruitbody data. In fact, the few higher taxa that were more speciose in the fruitbody data, were primarily non-soil taxa like *Polyporales, Hymenochaetales, Crepidotus*, etc. It was, however, interesting to note that *Ganoderma applanatum* (a wood decaying polypore) was found as OTU in 51 of 130 sites, including several sites with no trees, suggesting the species to be abundantly present in the spore bank. Although soil sampling catches fungi associated with above ground carbon sources, our results indicate that these are heavily undersampled. Studies indicate, that this part compose a major proportion of the total funga (Unterseher *et al*., 2011; Arnold and Lutzoni, 2007; Arnold, 2007), so to get a more complete estimate of the total fungal community, DNA-based methods will need to include sampling of above ground structures.

### 4.3 Detection of red-listed species

The fruitbody survey registered more red-listed species in total and average per site. However, when adjusting for red-listed species not present in the DNA reference database (476 of the 656 Danish red-listed species were present with sequence data in UNITE), and omitting red-listed species associated with dead wood and other non-soil resources, the eDNA-metabarcoding approach performed almost as good. However, the methods partly recorded different red-listed species, indicating that fruitbody surveys and eDNA-metabarcoding could be used complementarily to get a more reliable assessment of local conservation value, which to some degree conflicts with the findings of Runnel et al. (2015) that found fruitbody surveys to be superior to eDNA-based sampling of redlisted wood-inhabiting polypores at stand scale. The detection of red-listed species from environmental DNA samples must be expected to increase as sequence databases become more complete and well-annotated.

### 4.4 Species turnover comparable

Our results showed that community composition estimated from DNA-metabarcoding data correlated well with estimates based on fruitbody data. This correlation did not change much after narrowing taxonomic focus to Agaricomycetes/Agaricales, indicating that all approaches are suitable for describing fungal communities and species turnover along environmental gradients. However, eDNA-metabarcoding outperformed fruitbody data when it came to correlation with environmental gradients expressed by independent environmental variables for all subsets of data. Further, it appears that the wider soil fungal community is more predictable than the fruitbody community. This could be caused by fruitbodies constituting a more stochastic subset of the total funga, or alternatively that Agaricomycetes and Agaricales depend on less easily measured properties of the environment. Most species of Agaricomycetes and Agaricales produce billions of spores that are effectively dispersed (Peay & Bruns 2014) – which is not the case for some of the other main groups of fungi in this study (Money 2016). Hence the detected community of these by eDNA may in part be a signal from the spore bank. The spore bank community has been shown to have relatively low correlation with the active community of the same taxa for pine associated ectomycorrhizal fungi (Glassman et al 2015), and thus, the lower correlation seen in our study may potentially be caused by a similar discrepancy. The lower performance of fruitbody survey data likely also indicates that fruitbody formation is more sensitive to e.g. unpredictable variation in weather conditions.

As seen from the taxonomic composition the eDNA-metabarcoding has a much higher proportion of Ascomycota and other phyla of ‘micro-fungi’, but also a relatively lower proportion of non-soil Ascomycota and Basidiomycota. DNA-metabarcoding thus targets a community with a larger proportion of micro-fungi (possibly also due to PCR amplification biases), which must be assumed to be more dependent on soil composition and humidity, whereas the fruitbody data targets a community of macrofungi with a larger dependence on the vegetation and above ground conditions. This is reflected in light being among the best explanatory variables for the fruitbody data, and soil moisture and pH for the eDNA-metabarcoding data.

### 4.5 Sampling efficiency/depth

The results obtained in this study reflect the exact sampling protocols for bothfruitbody survey and eDNA sampling, as well as the bioinformatics processing of the sequence data. The fruitbody survey included three visits to each site, and it is obvious that more sampling visits continuously will add to the species list, and may be necessary to get a fully representative sample (Halme and Kotiaho, 2012; Newton *et al*., 2003; Straatsma *et al*., 2015) (but see Abrego et al. 2016). The (eDNA) soil sampling method included the mixing of 81 soil cores and thus several kilos of soil for each site, and was uniquely large compared to previous studies (e.g. Porter *et al*., 2008; Geml *et al*., 2010; Baptista *et al*., 2015; Pellissier *et al*., 2014; Geml *et al*., 2009; Schmidt *et al*., 2013). However, it still covered only 0.01 % of the soil surface of the 40 x 40 m sites, and of the approximately 5-20 kg soil sampled from each plot only 4 g of soil was used for DNA extraction. Also, we made no attempt to maximize coverage of visible variation at the sites but sampled completely systematically. Hence, both sampling approaches could be both up- and down-scaled for applications in practice. A study in Switzerland (Straatsma *et al*., 2015) recorded fruitbodies on a weekly basis over 21 years, and identified 101 species on average per year (408 species in total) in a forest study area close to ours in size (1,500 vs 1,600 m^2^). Although, their total number exceeds the site average of 68 fruitbody species (and 236 OTUs) in our study, their yearly average of 101 is only slightly higher than the average (93.5) of our forest/plantation sites after three 1 h visits, and we predict that it would require much further effort to get a significantly larger average species number for the fruitbody data. Presently, there is little knowledge on which parameters are most important for getting a representative sample with the eDNA-metabarcoding approach. We expect that extracting and sequencing the 81 soil cores separately or sequencing many sub-samples of the bulk sample would increase the number of detected OTUs, but this would also pose a marked increase of lab consumables and processing time. As many fungal mycelia must be assumed to be restricted in size and/or time, and as our bulk sample only covers 0.01 % of the soil surface, we expect that additional bulk samples – done at the same time or at another time of the year – would be the most cost-efficient way of capturing a larger sample of the real fungal community.

### 4.6 Practical applications

Both approaches represent surveys that are realistic to perform within the limits of standard surveys and research studies, and expenses were roughly comparable. For the full data, eDNA-metabarcoding resulted in more species (OTU) observations (30,668) than the fruitbody survey (8,793), whereas the numbers were more similar for Agaricomycetes (9,091 vs. 7,214) and Agaricales (4,507 vs. 4,334). In our data, fruitbody richness was a relatively weak predictor of total fungal richness as assessed with DNA metabarcoding, but was a relatively good predictor of the richness of fruitbody forming fungi (Agaricomycetes and Agaricales). This indicates that species richness of these groups may be assessed interchangeably with eDNA-metabarcoding or as fruitbodies.

For detection of red-listed species, eDNA-metabarcoding performed much better than expected, but still we would recommend a manual search for fruitbodies in all cases where larger areas need to be surveyed e.g. for conservation value, as also suggested for wood-inhabiting fungi (Runnel *et al*., 2015). However, eDNA-metabarcoding may supply valuable information, in cases of poor fruiting conditions, or in more targeted plot-based monitoring programmes. Fruitbody and eDNA-metabarcoding data result in comparable measures of species turnover, and thus, our results indicate that data may be combined for example to evaluate long time series including historical fruitbody data and future DNA-based surveys.

## Supporting information

## Acknowledgements

Karl-Henrik Larsson, Leif Örstadius and Viacheslav Spirin are thanked for identification of critical fungal collections, Anne Storgaard, Annegrete Eriksen, David Boertmann, Erik Arnfred Thomsen, Hanne Petra Katballe, Jette Anitha Hansen, Leif Wegge Laursen, Thomas Kehlet, Tom Smidth, Torbjørn Borgen are thanked for assistance in the field during fruitbody inventories. JHC acknowledge the Danish National Research Foundation for funding for the Center for Macroecology, Evolution and Climate, grant no. DNRF96.

## Conflicts of interest

All authors declare no conflicting interests.

## Authors’ contributions

All authors designed the study, TL collected the fruitbody data, TGF collected and analyzed the molecular data, TGF carried out the statistical analyses, TGF, JHC and RK wrote first draft, all authors contributed to revising the paper.

## Data Accessibility

Sequence data is available from DataDryad, and files documenting the analyses from GitHub. Fruitbody data is deposited in GBIF. [[more information will be added before publication]].

