## Supplementary material for "Man against machine: Do fungal fruitbodies and eDNA give similar biodiversity assessments across broad environmental gradients?"

Supplementary methods.
This study was part of the Biowide project, and many aspects are presented and discussed in more detail in Brunbjerg et al. (2017).

Environmental variables.
Soil samples (0-10 cm, 5 cm diameter) were collected within 4 subplots of the 130 sites and separated in organic (Oa) and mineral (A/B) soil horizons. Across all sites, a total of 664 soil samples were collected. Organic horizons were separated from the mineral horizons when both were present. Soil pH was measured on 10g soil in 30 ml deionized water, shaken vigorously for 20 seconds, and then settling for 30 minutes. Measurements were done with a Mettler Toledo Seven Compact pH meter. Soil pH of the 0-10 cm soil layer was calculated weighted for the proportion of organic matter to mineral soil (average of samples taken in 4 subplots). Organic matter content was measured as the percentage of the 0-10 cm core that was organic matter. 129 of the total samples were measured for carbon content (LECO elemental analyzer) and total phosphorus content (H2SO4-Se digestion and colorimetric analysis). NIR was used to analyze each sample for total carbon and phosphorus concentrations. Reflectance spectra was analyzed within a range of 10000-4000 cm-1 with a Antaris II NIR spectrophotometer (Thermo Fisher Scientific). A partial least square regression was used to test for a correlation between the NIR data and the subset reference analyses to calculate total carbon and phosphorous (see Brunbjerg et al. 2017 for more details).

Supplementary Figures

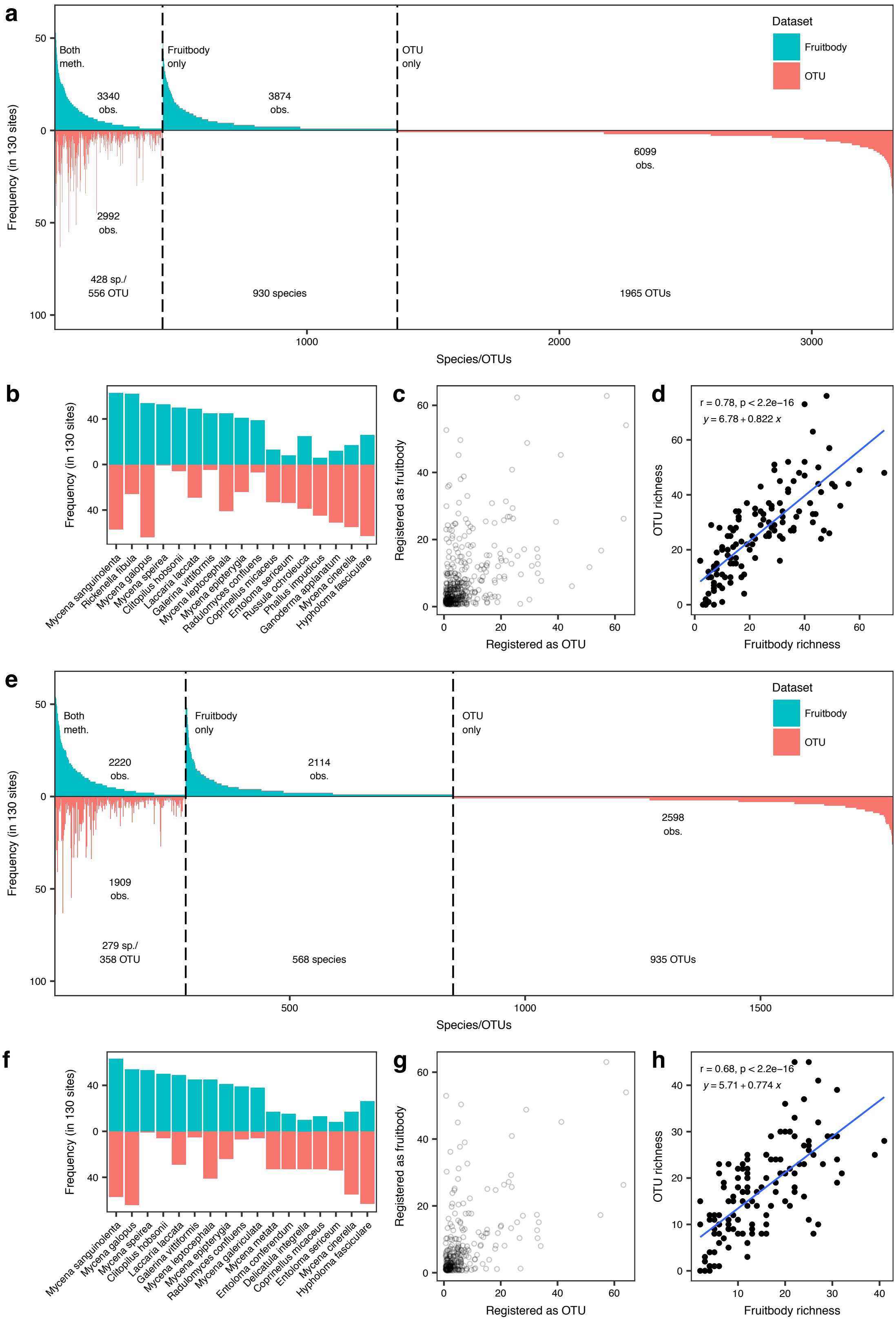

Supplementary Figure 1. Frequency of species and OTUs among the 130 sampling sites, restricted to Agaricomycetes (a-e) and Agaricales (f-h). a) Frequency of Agaricomycetes species and OTUs sorted by decreasing frequency, and grouped by species/OTUs recorded with both methods or only by either DNA or as fruitbody, y-axis indicates the number of sampling sites (of 130) in which a species or OTU was found, number of species/OTUs and number of observations are indicated for each group. b) Top 10 most frequent Agaricomycetes species recorded with either method. c) scatterplot of fruitbody-based frequency vs DNA frequency of the 428 Agaricomycetes species recorded by both methods. d) Species/OTU richness of the 130 sites as recorded with fruitbody or DNA for the Agaricomycetes 428 species recorded with both methods. e) Frequency of Agaricales species and OTUs sorted by decreasing frequency, and grouped by species/OTUs recorded with both methods or only by either DNA or as fruitbody, y-axis indicates the number of sampling sites (of 130) in which a species or OTU was found, number of species/OTUs and number of observations are indicated for each group. f) Top 10 most frequent Agaricales species recorded as with either method. g) scatterplot of fruitbody-based frequency vs DNA frequency of the 279 Agaricales species recorded by both methods. h) Species/OTU richness of the 130 sites as recorded with fruitbody or DNA for the 279 Agaricales species recorded with both methods.

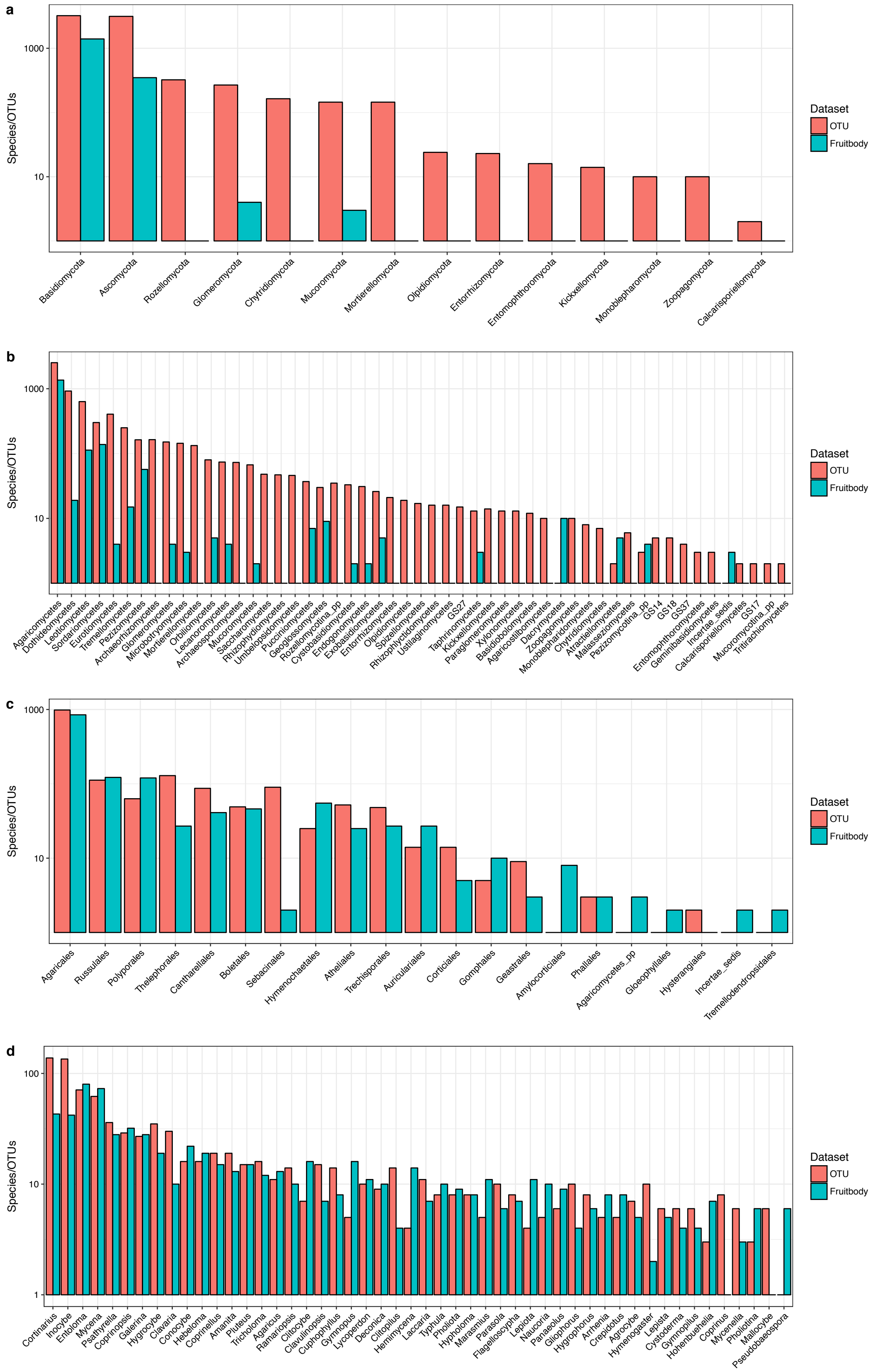

Supplementary Figure 2. Taxonomic composition of DNA metabarcoding and fruitbody data. a) Number of species in each phylum. b) Number of species in each class. c) Number of species in each order of Agaricomycetes. d) Number of species in each genus of Agaricales (restricted to genera with at least 5 species recorded). Y-axis is logarithmic.

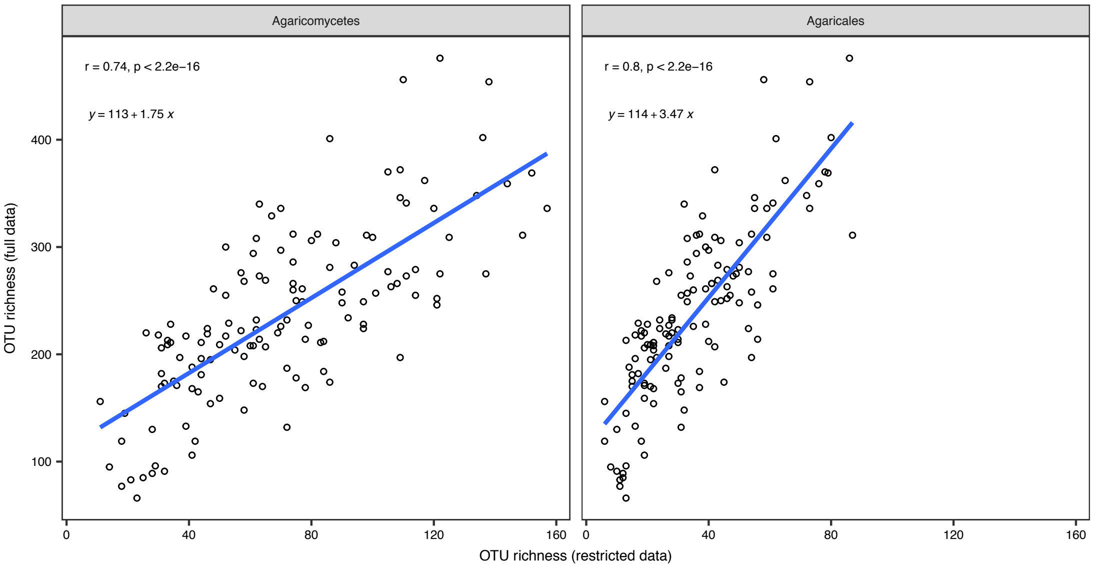

Supplementary figure 3. Correlation between OTU richness based on taxonomic subsets. Blue lines represent the linear regression of OTU richness of the taxonomically restricted dataset against OTU richness based on the full data, while the dotted line shows the identity line (x=y). Correlations are shown for the taxonomic subsets Agaricomycetes and Agaricales).

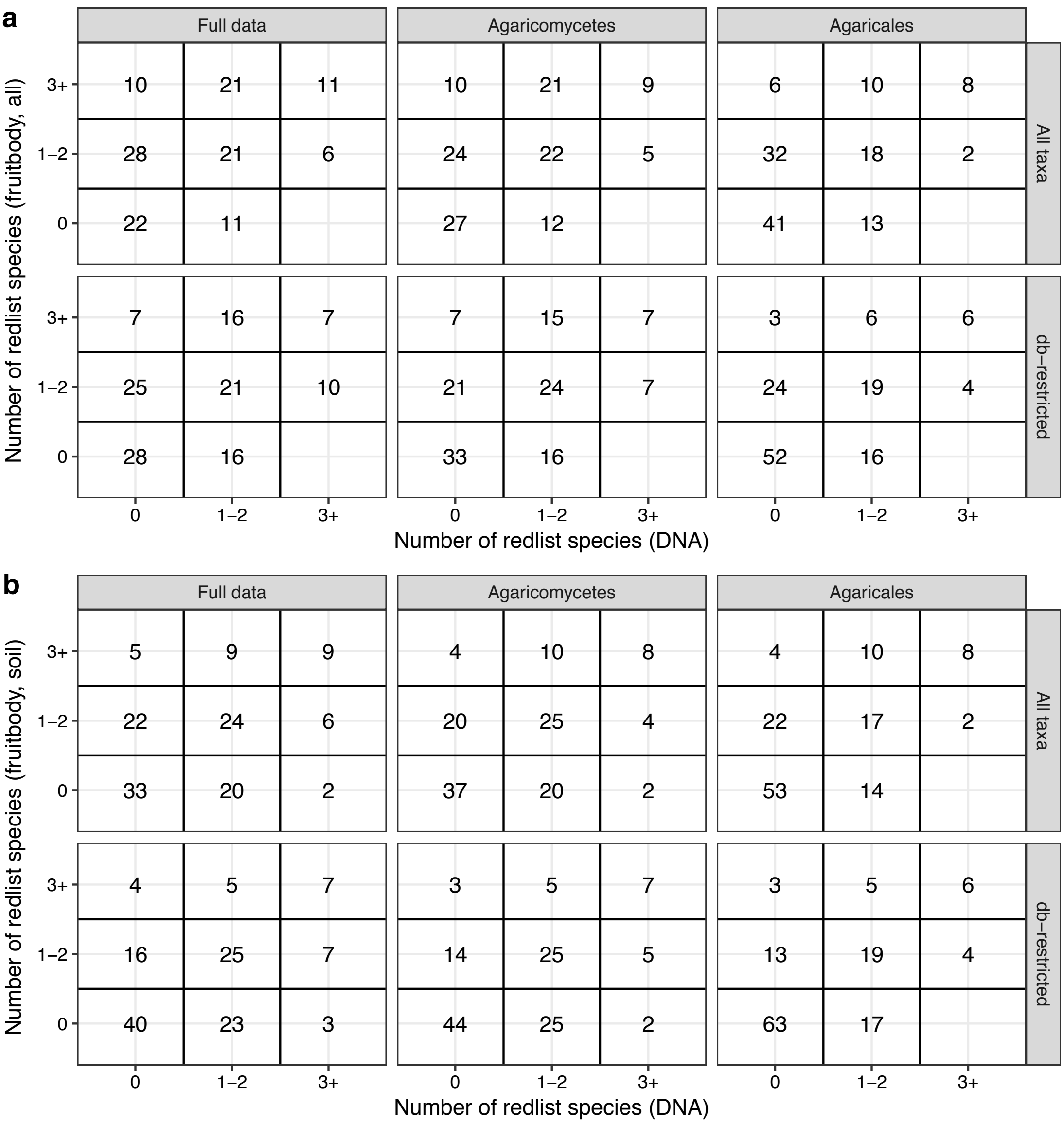

Supplementary figure 4. Red-listed species. Number of red-listed species recorded at each site with either method. Upper panel shows data for all species, lower panel is restricted to red-listed species present in the DNA reference database (and thus possible to identify with both methods), a) shows data for the full fruitbody dataset, b) shows data for soil fruitbody data only.

### Supplementary Tables

#### Supplementary table 1. Number of species per phylum

| Phylum | OTU | Fruitbody total | Soil frb. | Non-soil frb. |
|---|---|---|---|---|
| Ascomycota | 3143 (39 %) | 348 (20 %) | 105 (10 %) | 243 (36 %) |
| Basidiomycota | 3221 (40 %) | 1398 (80 %) | 958 (90 %) | 440 (64 %) |
| Calcarisporiellomycota | 1 (0 %) | - | - | - |
| Chytridiomycota | 163 (2 %) | - | - | - |
| Entomophthoromycota | 15 (0 %) | - | - | - |
| Entorrhizomycota | 22 (0 %) | - | - | - |
| Glomeromycota | 267 (3 %) | 3 (0 %) | 3 (0 %) | - |
| Kickxellomycota | 13 (0 %) | - | - | - |
| Monoblepharomycota | 9 (0 %) | - | - | - |
| Mortierellomycota | 144 (2 %) | - | - | - |
| Mucoromycota | 144 (2 %) | 2 (0 %) | 1 (0 %) | 1 (0 %) |
| Olpidiomycota | 23 (0 %) | - | - | - |
| Rozellomycota | 322 (4 %) | - | - | - |
| unidentified | 614 (8 %) | - | - | - |
| Zoopagomycota | 9 (0 %) | - | - | - |

Supplementary table 2. Number of species per class

| Class | OTU | Fruitbody total | Soil frb. | Non-soil frb. |
|---|---|---|---|---|
| Agaricomycetes | 2521 (31 %) | 1358 (78 %) | 955 (90 %) | 403 (59 %) |
| Agaricostilbomycetes | 9 (0 %) | - | - | - |
| Archaeorhizomycetes | 163 (2 %) | - | - | - |
| Archaeosporomycetes | 72 (1 %) | - | - | - |
| Atractiellomycetes | 1 (0 %) | 4 (0 %) | - | 4 (1 %) |
| Basidiobolomycetes | 11 (0 %) | - | - | - |
| Calcarisporiellomycetes | 1 (0 %) | - | - | - |
| Chytridiomycetes | 6 (0 %) | - | - | - |
| Cystobasidiomycetes | 32 (0 %) | 1 (0 %) | - | 1 (0 %) |
| Dacrymycetes | - | 9 (1 %) | 1 (0 %) | 8 (1 %) |
| Dothideomycetes | 920 (11 %) | 18 (1 %) | 1 (0 %) | 17 (2 %) |
| Endogonomycetes | 30 (0 %) | 1 (0 %) | 1 (0 %) | - |
| Entomophthoromycetes | 2 (0 %) | - | - | - |
| Entorrhizomycetes | 20 (0 %) | - | - | - |
| Eurotiomycetes | 404 (5 %) | 3 (0 %) | 2 (0 %) | 1 (0 %) |
| Exobasidiomycetes | 25 (0 %) | 4 (0 %) | - | 4 (1 %) |
| Geminibasidiomycetes | 2 (0 %) | - | - | - |
| Geoglossomycetes | 29 (0 %) | 8 (0 %) | 8 (1 %) | - |
| Glomeromycetes | 150 (2 %) | 3 (0 %) | 3 (0 %) | - |
| GS14 | 4 (0 %) | - | - | - |
| GS17 | 1 (0 %) | - | - | - |
| GS18 | 4 (0 %) | - | - | - |
| GS27 | 14 (0 %) | - | - | - |
| GS37 | 3 (0 %) | - | - | - |
| Incertae_sedis | - | 2 (0 %) | - | 2 (0 %) |
| Kickxellomycetes | 13 (0 %) | - | - | - |
| Lecanoromycetes | 73 (1 %) | 3 (0 %) | - | 3 (0 %) |
| Leotiomycetes | 632 (8 %) | 112 (6 %) | 22 (2 %) | 90 (13 %) |
| Malasseziomycetes | 5 (0 %) | - | - | - |
| Microbotryomycetes | 143 (2 %) | 2 (0 %) | - | 2 (0 %) |
| Monoblepharidomycetes | 7 (0 %) | - | - | - |
| Mortierellomycetes | 132 (2 %) | - | - | - |
| Mucoromycetes | 66 (1 %) | 1 (0 %) | - | 1 (0 %) |
| Mucoromycotina__pp | 1 (0 %) | - | - | - |
| Olpidiomycetes | 18 (0 %) | - | - | - |
| Orbiliomycetes | 79 (1 %) | 4 (0 %) | - | 4 (1 %) |
| Paraglomeromycetes | 12 (0 %) | - | - | - |
| Pezizomycetes | 162 (2 %) | 56 (3 %) | 50 (5 %) | 6 (1 %) |
| Pezizomycotina__pp | 2 (0 %) | 3 (0 %) | - | 3 (0 %) |
| Pucciniomycetes | 36 (0 %) | 6 (0 %) | 1 (0 %) | 5 (1 %) |

| | | | | |
|---|---|---|---|---|
| Rhizophlyctidomycetes | 15 (0 %) | - | - | - |
| Rhizophydiomycetes | 46 (1 %) | - | - | - |
| Rozellomycotina__pp | 34 (0 %) | - | - | - |
| Saccharomycetes | 47 (1 %) | - | - | - |
| Sordariomycetes | 300 (4 %) | 137 (8 %) | 22 (2 %) | 115 (17 %) |
| Spizellomycetes | 16 (0 %) | - | - | - |
| Taphrinomycetes | 12 (0 %) | 2 (0 %) | - | 2 (0 %) |
| Tremellomycetes | 249 (3 %) | 14 (1 %) | 1 (0 %) | 13 (2 %) |
| Tritirachiomycetes | 1 (0 %) | - | - | - |
| Umbelopsidomycetes | 45 (1 %) | - | - | - |
| unidentified | 1504 (19 %) | - | - | - |
| Ustilaginomycetes | 15 (0 %) | - | - | - |
| Xylonomycetes | 12 (0 %) | - | - | - |
| Zoopagomycetes | 9 (0 %) | - | - | - |

Supplementary table 3. Number of species per order in Agaricomycetes

| Order | OTU | Fruitbody total | Soil frb. | Non-soil frb. |
|---|---|---|---|---|
| Agaricales | 1293 (51 %) | 847 (62 %) | 710 (74 %) | 137 (34 %) |
| Agaricomycetes_pp | - | 2 (0 %) | - | 2 (0 %) |
| Amylocorticiales | - | 7 (1 %) | 1 (0 %) | 6 (1 %) |
| Atheliales | 60 (2 %) | 24 (2 %) | 7 (1 %) | 17 (4 %) |
| Auriculariales | 78 (3 %) | 26 (2 %) | - | 26 (6 %) |
| Boletales | 52 (2 %) | 45 (3 %) | 42 (4 %) | 3 (1 %) |
| Cantharellales | 170 (7 %) | 40 (3 %) | 23 (2 %) | 17 (4 %) |
| Corticiales | 21 (1 %) | 4 (0 %) | - | 4 (1 %) |
| Geastrales | 12 (0 %) | 2 (0 %) | 2 (0 %) | - |
| Gloeophyllales | - | 1 (0 %) | - | 1 (0 %) |
| Gomphales | 4 (0 %) | 9 (1 %) | 6 (1 %) | 3 (1 %) |
| GS28 | 1 (0 %) | - | - | - |
| Hymenochaetales | 36 (1 %) | 54 (4 %) | 5 (1 %) | 49 (12 %) |
| Hysterangiales | 1 (0 %) | - | - | - |
| Incertae_sedis | - | 1 (0 %) | - | 1 (0 %) |
| Phallales | 2 (0 %) | 2 (0 %) | 2 (0 %) | - |
| Polyporales | 75 (3 %) | 119 (9 %) | 25 (3 %) | 94 (23 %) |
| Russulales | 117 (5 %) | 121 (9 %) | 86 (9 %) | 35 (9 %) |
| Sebacinales | 205 (8 %) | 1 (0 %) | 1 (0 %) | - |
| Thelephorales | 167 (7 %) | 26 (2 %) | 26 (3 %) | - |
| Trechisporales | 79 (3 %) | 26 (2 %) | 18 (2 %) | 8 (2 %) |
| Tremellodendropsidales | 7 (0 %) | 1 (0 %) | 1 (0 %) | - |
| unidentified | 141 (6 %) | - | - | - |

Supplementary table 4. Number of species per genus in Agaricales

| Genus | OTU | Fruitbody total | Soil frb. | Non-soil frb. |
|---|---|---|---|---|
| Agaricus | 10 (1 %) | 12 (1 %) | 12 (2 %) | - |
| Agrocybe | 6 (0 %) | 4 (0 %) | 4 (1 %) | - |
| Alboleptonia | 1 (0 %) | - | - | - |
| Alnicola | 3 (0 %) | - | - | - |
| Amanita | 18 (1 %) | 12 (1 %) | 12 (2 %) | - |
| Ampulloclitocybe | 1 (0 %) | 1 (0 %) | 1 (0 %) | - |
| Aphanobasidium | 1 (0 %) | 3 (0 %) | - | 3 (2 %) |
| Armillaria | - | 3 (0 %) | 3 (0 %) | - |
| Arrhenia | 4 (0 %) | 7 (1 %) | 7 (1 %) | - |
| Aspropaxillus | 1 (0 %) | 1 (0 %) | 1 (0 %) | - |
| Asterophora | 1 (0 %) | 2 (0 %) | 2 (0 %) | - |
| Atheniella | 1 (0 %) | 2 (0 %) | 2 (0 %) | - |
| Baeospora | 1 (0 %) | 1 (0 %) | 1 (0 %) | - |
| Bolbitius | 1 (0 %) | 2 (0 %) | 2 (0 %) | - |
| Bovista | 2 (0 %) | 4 (0 %) | 4 (1 %) | - |
| Calocybe | - | 1 (0 %) | 1 (0 %) | - |
| Calyptella | 3 (0 %) | 4 (0 %) | - | 4 (3 %) |
| Camarophyllopsis | 4 (0 %) | 2 (0 %) | 2 (0 %) | - |
| Cellypha | - | 1 (0 %) | - | 1 (1 %) |
| Cephaloscypha | - | 1 (0 %) | - | 1 (1 %) |
| Chlorophyllum | - | 2 (0 %) | 2 (0 %) | - |
| Chondrostereum | - | 1 (0 %) | - | 1 (1 %) |
| Chromocyphella | - | 1 (0 %) | - | 1 (1 %) |
| Cibaomyces | 1 (0 %) | - | - | - |
| Clavaria | 29 (2 %) | 9 (1 %) | 9 (1 %) | - |
| Clavulinopsis | 14 (1 %) | 6 (1 %) | 6 (1 %) | - |
| Clitocella | - | 1 (0 %) | 1 (0 %) | - |
| Clitocybe | 6 (0 %) | 15 (2 %) | 15 (2 %) | - |
| Clitopilus | 13 (1 %) | 3 (0 %) | 3 (0 %) | - |
| Collybia | - | 3 (0 %) | 3 (0 %) | - |
| Conocybe | 15 (1 %) | 21 (2 %) | 21 (3 %) | - |
| Coprinellus | 18 (1 %) | 14 (2 %) | 14 (2 %) | - |
| Coprinopsis | 28 (2 %) | 31 (4 %) | 31 (4 %) | - |
| Coprinus | 7 (1 %) | - | - | - |
| Cortinarius | 137 (11 %) | 42 (5 %) | 42 (6 %) | - |
| Crepidotus | 4 (0 %) | 7 (1 %) | - | 7 (5 %) |
| Crinipellis | 1 (0 %) | 1 (0 %) | 1 (0 %) | - |
| Cristinia | 2 (0 %) | 3 (0 %) | 3 (0 %) | - |
| Cuphophyllus | 13 (1 %) | 7 (1 %) | 7 (1 %) | - |
| Cyathus | 1 (0 %) | 1 (0 %) | 1 (0 %) | - |

| | | | | |
|---|---|---|---|---|
| Cyclocybe | 1 (0 %) | - | - | - |
| Cylindrobasidium | 1 (0 %) | 1 (0 %) | - | 1 (1 %) |
| Cystoderma | 5 (0 %) | 3 (0 %) | 3 (0 %) | - |
| Cystodermella | - | 1 (0 %) | 1 (0 %) | - |
| Cystolepiota | - | 2 (0 %) | 2 (0 %) | - |
| Deconica | 8 (1 %) | 9 (1 %) | 6 (1 %) | 3 (2 %) |
| Delicatula | 1 (0 %) | 1 (0 %) | 1 (0 %) | - |
| Dendrothele | - | 2 (0 %) | - | 2 (1 %) |
| Dermoloma | - | 2 (0 %) | 2 (0 %) | - |
| Disciseda | - | 1 (0 %) | 1 (0 %) | - |
| Entocybe | 2 (0 %) | - | - | - |
| Entoloma | 70 (5 %) | 79 (9 %) | 79 (11 %) | - |
| Eonema | - | 1 (0 %) | - | 1 (1 %) |
| Episphaeria | - | 1 (0 %) | - | 1 (1 %) |
| Fayodia | - | 1 (0 %) | 1 (0 %) | - |
| Fistulina | - | 1 (0 %) | - | 1 (1 %) |
| Flagelloscypha | 7 (1 %) | 6 (1 %) | - | 6 (4 %) |
| Flammula | - | 2 (0 %) | - | 2 (1 %) |
| Flammulaster | 1 (0 %) | 4 (0 %) | 3 (0 %) | 1 (1 %) |
| Flammulina | - | 2 (0 %) | - | 2 (1 %) |
| Galerina | 26 (2 %) | 27 (3 %) | 25 (4 %) | 2 (1 %) |
| Gliophorus | 9 (1 %) | 3 (0 %) | 3 (0 %) | - |
| Globulicium | 1 (0 %) | - | - | - |
| Gloiocephala | - | 1 (0 %) | - | 1 (1 %) |
| Gloioxanthomyces | 1 (0 %) | 1 (0 %) | 1 (0 %) | - |
| Gymnopilus | 5 (0 %) | 3 (0 %) | 2 (0 %) | 1 (1 %) |
| Gymnopus | 4 (0 %) | 15 (2 %) | 13 (2 %) | 2 (1 %) |
| Hebeloma | 15 (1 %) | 18 (2 %) | 18 (3 %) | - |
| Hemimycena | 3 (0 %) | 13 (2 %) | 5 (1 %) | 8 (6 %) |
| Henningsomyces | - | 2 (0 %) | - | 2 (1 %) |
| Hodophilus | 3 (0 %) | 2 (0 %) | 2 (0 %) | - |
| Hohenbuehelia | 2 (0 %) | 6 (1 %) | 2 (0 %) | 4 (3 %) |
| Homophron | 1 (0 %) | 1 (0 %) | 1 (0 %) | - |
| Hydropus | 3 (0 %) | 2 (0 %) | 2 (0 %) | - |
| Hygrocybe | 34 (3 %) | 18 (2 %) | 18 (3 %) | - |
| Hygrophorus | 7 (1 %) | 5 (1 %) | 5 (1 %) | - |
| Hymenogaster | 9 (1 %) | 1 (0 %) | 1 (0 %) | - |
| Hymenopellis | 2 (0 %) | 1 (0 %) | 1 (0 %) | - |
| Hyphodontiella | 3 (0 %) | 1 (0 %) | - | 1 (1 %) |
| Hypholoma | 7 (1 %) | 7 (1 %) | 5 (1 %) | 2 (1 %) |
| Infundibulicybe | 1 (0 %) | 1 (0 %) | 1 (0 %) | - |
| Inocybe | 134 (10 %) | 41 (5 %) | 41 (6 %) | - |

| | | | | |
|---|---|---|---|---|
| Kuehneromyces | 1 (0 %) | 1 (0 %) | - | 1 (1 %) |
| Laccaria | 10 (1 %) | 6 (1 %) | 6 (1 %) | - |
| Lachnella | - | 2 (0 %) | - | 2 (1 %) |
| Lacrymaria | 3 (0 %) | 2 (0 %) | 2 (0 %) | - |
| Lepiota | 3 (0 %) | 10 (1 %) | 10 (1 %) | - |
| Lepista | 5 (0 %) | 4 (0 %) | 4 (1 %) | - |
| Leratiomyces | - | 1 (0 %) | 1 (0 %) | - |
| Leucoagaricus | 2 (0 %) | 2 (0 %) | 2 (0 %) | - |
| Leucocybe | - | 1 (0 %) | 1 (0 %) | - |
| Lichenomphalia | - | 1 (0 %) | 1 (0 %) | - |
| Lindtneria | 3 (0 %) | 3 (0 %) | 3 (0 %) | - |
| Lycoperdon | 9 (1 %) | 10 (1 %) | 10 (1 %) | - |
| Lyophyllum | 2 (0 %) | 2 (0 %) | 2 (0 %) | - |
| Macrocystidia | 1 (0 %) | 1 (0 %) | 1 (0 %) | - |
| Macrolepiota | 1 (0 %) | 4 (0 %) | 4 (1 %) | - |
| Macrotyphula | 1 (0 %) | 2 (0 %) | 2 (0 %) | - |
| Maireina | - | 2 (0 %) | - | 2 (1 %) |
| Mallocybe | 5 (0 %) | - | - | - |
| Marasmiellus | - | 2 (0 %) | - | 2 (1 %) |
| Marasmius | 4 (0 %) | 10 (1 %) | 10 (1 %) | - |
| Megacollybia | 1 (0 %) | 1 (0 %) | 1 (0 %) | - |
| Melanoleuca | 1 (0 %) | 3 (0 %) | 3 (0 %) | - |
| Melanophyllum | - | 1 (0 %) | 1 (0 %) | - |
| Merismodes | - | 2 (0 %) | - | 2 (1 %) |
| Merulicium | - | 1 (0 %) | 1 (0 %) | - |
| Mucidula | 1 (0 %) | 1 (0 %) | - | 1 (1 %) |
| Mucronella | - | 3 (0 %) | - | 3 (2 %) |
| Mycena | 61 (5 %) | 72 (9 %) | 51 (7 %) | 21 (15 %) |
| Mycenastrum | 1 (0 %) | - | - | - |
| Mycenella | 5 (0 %) | 2 (0 %) | 2 (0 %) | - |
| Mycetinis | 3 (0 %) | 3 (0 %) | 3 (0 %) | - |
| Mycocalia | - | 2 (0 %) | 2 (0 %) | - |
| Myochromella | 1 (0 %) | 1 (0 %) | 1 (0 %) | - |
| Naucoria | 4 (0 %) | 9 (1 %) | 9 (1 %) | - |
| Nematoctonus | 1 (0 %) | - | - | - |
| Neohygrocybe | 2 (0 %) | - | - | - |
| Omphaliaster | - | 1 (0 %) | 1 (0 %) | - |
| Pachylepyrium | 1 (0 %) | - | - | - |
| Panaeolus | 5 (0 %) | 8 (1 %) | 8 (1 %) | - |
| Panellus | - | 2 (0 %) | - | 2 (1 %) |
| Paralepista | 1 (0 %) | 2 (0 %) | 2 (0 %) | - |
| Parasola | 9 (1 %) | 5 (1 %) | 5 (1 %) | - |

| | | | | |
|---|---|---|---|---|
| Phaeocollybia | 1 (0 %) | - | - | - |
| Phaeogalera | 1 (0 %) | - | - | - |
| Phaeolepiota | 1 (0 %) | - | - | - |
| Phaeomarasmius | - | 1 (0 %) | - | 1 (1 %) |
| Phaeonematoloma | 1 (0 %) | 1 (0 %) | 1 (0 %) | - |
| Phloeomana | 2 (0 %) | - | - | - |
| Pholiota | 7 (1 %) | 8 (1 %) | 5 (1 %) | 3 (2 %) |
| Pholiotina | 2 (0 %) | 5 (1 %) | 5 (1 %) | - |
| Pistillina | - | 1 (0 %) | - | 1 (1 %) |
| Plicatura | - | 1 (0 %) | - | 1 (1 %) |
| Pluteus | 14 (1 %) | 14 (2 %) | 3 (0 %) | 11 (8 %) |
| Porotheleum | - | 1 (0 %) | - | 1 (1 %) |
| Protoglossum | 2 (0 %) | - | - | - |
| Protostropharia | - | 1 (0 %) | 1 (0 %) | - |
| Psathyrella | 35 (3 %) | 27 (3 %) | 24 (3 %) | 3 (2 %) |
| Pseudobaeospora | - | 5 (1 %) | 5 (1 %) | - |
| Pseudoclitocybe | 1 (0 %) | 1 (0 %) | 1 (0 %) | - |
| Pseudolasiobolus | - | 1 (0 %) | - | 1 (1 %) |
| Pseudotricholoma | - | 1 (0 %) | 1 (0 %) | - |
| Psilocybe | 2 (0 %) | 4 (0 %) | 4 (1 %) | - |
| Pterula | - | 2 (0 %) | 2 (0 %) | - |
| Radulomyces | 1 (0 %) | 1 (0 %) | - | 1 (1 %) |
| Ramariopsis | 13 (1 %) | 9 (1 %) | 9 (1 %) | - |
| Rectipilus | 1 (0 %) | 1 (0 %) | - | 1 (1 %) |
| Resinomycena | - | 1 (0 %) | - | 1 (1 %) |
| Resupinatus | - | 4 (0 %) | - | 4 (3 %) |
| Rhizomarasmius | - | 1 (0 %) | 1 (0 %) | - |
| Rhodocollybia | 1 (0 %) | 2 (0 %) | 2 (0 %) | - |
| Rhodocybe | - | 2 (0 %) | 2 (0 %) | - |
| Rhodophana | - | 1 (0 %) | 1 (0 %) | - |
| Rimbachia | - | 2 (0 %) | 2 (0 %) | - |
| Ripartites | 2 (0 %) | 1 (0 %) | 1 (0 %) | - |
| Roridomyces | 1 (0 %) | 1 (0 %) | 1 (0 %) | - |
| Rugosomyces | - | 1 (0 %) | 1 (0 %) | - |
| Sagaranella | 1 (0 %) | 1 (0 %) | 1 (0 %) | - |
| Sarcomyxa | - | 1 (0 %) | - | 1 (1 %) |
| Schizophyllum | - | 1 (0 %) | - | 1 (1 %) |
| Seticyphella | - | 2 (0 %) | - | 2 (1 %) |
| Simocybe | - | 4 (0 %) | 1 (0 %) | 3 (2 %) |
| Sphagnurus | 1 (0 %) | 1 (0 %) | 1 (0 %) | - |
| Stephanospora | 1 (0 %) | - | - | - |
| Strobilurus | - | 1 (0 %) | 1 (0 %) | - |

| | | | | |
|---|---|---|---|---|
| Stropharia | 2 (0 %) | 4 (0 %) | 4 (1 %) | - |
| Tephrocybe | 1 (0 %) | - | - | - |
| Tricholoma | 15 (1 %) | 11 (1 %) | 11 (2 %) | - |
| Tricholomopsis | 2 (0 %) | 1 (0 %) | - | 1 (1 %) |
| Tubaria | 4 (0 %) | 4 (0 %) | 4 (1 %) | - |
| Typhrasa | - | 1 (0 %) | 1 (0 %) | - |
| Typhula | 7 (1 %) | 9 (1 %) | 6 (1 %) | 3 (2 %) |
| unidentified | 310 (24 %) | - | - | - |
| Volvopluteus | 1 (0 %) | - | - | - |
| Xerula | - | 1 (0 %) | 1 (0 %) | - |

Supplementary table 5. Red-listed species. Frequency of registration as fruitbody/OTU across 130 sites, whether fruitbody species was recorded as 'soil species', whether the species was present in the DNA reference database (Ref-db), the taxonomic affiliation (Order, Class), and the Species Hypothesis code (SH) according to UNITE database.

| Species | Fruitbody | OTU | Soil | Ref-db | Order | Class | SH |
|---|---|---|---|---|---|---|---|
| Agaricus porphyrizon | 1 | | x | | Agaricales | Agaricomycetes | |
| Agrocybe firma | | 1 | | | Agaricales | Agaricomycetes | SH001883.07FU |
| Agrocybe vervacti | 1 | | x | x | Agaricales | Agaricomycetes | |
| Amanita olivaceogrisea | 2 | 5 | x | x | Agaricales | Agaricomycetes | SH101622.07FU |
| Amanita strobiliformis | 1 | | x | x | Agaricales | Agaricomycetes | |
| Anomoloma myceliosum | 1 | | x | x | Amylocorticiales | Agaricomycetes | |
| Antrodia malicola | 1 | | | x | Polyporales | Agaricomycetes | |
| Arrhenia epichysium | 1 | | x | x | Agaricales | Agaricomycetes | |
| Arrhenia lobata | 1 | 1 | x | x | Agaricales | Agaricomycetes | SH196811.07FU |
| Arrhenia onisca | 2 | | x | | Agaricales | Agaricomycetes | |
| Aurantiporus croceus | 1 | | | x | Polyporales | Agaricomycetes | |
| Botryobasidium intertextum | 1 | | | x | Cantharellales | Agaricomycetes | |
| Buglossoporus quercinus | 1 | | | | Polyporales | Agaricomycetes | |
| Cabalodontia subcretacea | 1 | | | x | Polyporales | Agaricomycetes | |
| Camarophyllopsis schulzeri | 1 | 3 | x | x | Agaricales | Agaricomycetes | SH180406.07FU |
| Camarops tubulina | 1 | | | x | Boliniales | Sordariomycetes | |
| Ceriporia purpurea | 5 | | | x | Polyporales | Agaricomycetes | |
| Chromocyphella muscicola | 3 | | | | Agaricales | Agaricomycetes | |
| Clavaria flavipes | 2 | 4 | x | x | Agaricales | Agaricomycetes | SH185116.07FU |
| Clavaria incarnata | 2 | | x | | Agaricales | Agaricomycetes | |
| Clavaria tenuipes | | 3 | | | Agaricales | Agaricomycetes | SH197585.07FU |
| Clavicorona taxophila | 1 | | x | x | Russulales | Agaricomycetes | |
| Clavulinopsis microspora | 3 | | x | | Agaricales | Agaricomycetes | |
| Conocybe apala | | 1 | | | Agaricales | Agaricomycetes | SH179180.07FU (as Conocybe crispa) |
| Conocybe dumetorum | 4 | | x | x | Agaricales | Agaricomycetes | |
| Cortinarius bergeronii | 1 | 1 | x | x | Agaricales | Agaricomycetes | SH222487.07FU (as Cortinarius cedretorum) |
| Cortinarius catharinae | | 1 | | | Agaricales | Agaricomycetes | SH222352.07FU |
| Cortinarius cotoneus | | 1 | | | Agaricales | Agaricomycetes | SH222458.07FU |
| Cortinarius danicus | 1 | | x | | Agaricales | Agaricomycetes | |
| Cortinarius langeorum | 1 | | x | x | Agaricales | Agaricomycetes | |
| Cortinarius splendens | | 1 | | | Agaricales | Agaricomycetes | SH221481.07FU |
| Cuphophyllus colemannianus | 1 | 1 | x | x | Agaricales | Agaricomycetes | SH000585.07FU |
| Cuphophyllus flavipes | | 1 | | | Agaricales | Agaricomycetes | SH023983.07FU |
| Cuphophyllus fornicatus | 1 | | x | x | Agaricales | Agaricomycetes | |
| Cuphophyllus lacmus | | 3 | | | Agaricales | Agaricomycetes | SH020555.07FU |

| | | | | | | | |
|---|---|---|---|---|---|---|---|
| Cuphophyllus russocoriaceus | 2 | | x | | Agaricales | Agaricomycetes | |
| Dermoloma pseudocuneifolium | 3 | | x | | Agaricales | Agaricomycetes | |
| Disciseda bovista | 1 | | x | | Agaricales | Agaricomycetes | |
| Eichleriella deglubens | 5 | | | x | Auriculariales | Agaricomycetes | |
| Entoloma ameides | 1 | 2 | x | x | Agaricales | Agaricomycetes | SH197363.07FU (as Entoloma sacchariolens) |
| Entoloma anatinum | 1 | | x | x | Agaricales | Agaricomycetes | |
| Entoloma asprellum | | 1 | | | Agaricales | Agaricomycetes | SH000812.07FU |
| Entoloma atrocoeruleum | 2 | | x | x | Agaricales | Agaricomycetes | |
| Entoloma byssisedum | 3 | | x | x | Agaricales | Agaricomycetes | |
| Entoloma caesiocinctum | 1 | | x | x | Agaricales | Agaricomycetes | |
| Entoloma chalybaeum | 5 | | x | | Agaricales | Agaricomycetes | |
| Entoloma clandestinum | 5 | 6 | x | x | Agaricales | Agaricomycetes | SH189311.07FU |
| Entoloma corvinum | 4 | | x | | Agaricales | Agaricomycetes | |
| Entoloma cuspidiferum | | 1 | | | Agaricales | Agaricomycetes | SH189316.07FU |
| Entoloma dichroum | 1 | | x | | Agaricales | Agaricomycetes | |
| Entoloma elodes | 7 | | x | | Agaricales | Agaricomycetes | |
| Entoloma euchroum | 1 | | x | x | Agaricales | Agaricomycetes | |
| Entoloma exile | 7 | | x | x | Agaricales | Agaricomycetes | |
| Entoloma formosum | 5 | 2 | x | | Agaricales | Agaricomycetes | SH221238.07FU |
| Entoloma glaucobasis | 1 | 2 | x | x | Agaricales | Agaricomycetes | SH526267.07FU |
| Entoloma griseocyaneum | 4 | | x | x | Agaricales | Agaricomycetes | |
| Entoloma hirtum | 1 | | x | | Agaricales | Agaricomycetes | |
| Entoloma hispidulum | 1 | | x | | Agaricales | Agaricomycetes | |
| Entoloma incanum | 2 | 3 | x | x | Agaricales | Agaricomycetes | SH221239.07FU |
| Entoloma infula | 3 | | x | x | Agaricales | Agaricomycetes | |
| Entoloma lepidissimum | 1 | | x | x | Agaricales | Agaricomycetes | |
| Entoloma lividocyanulum | 1 | | x | | Agaricales | Agaricomycetes | |
| Entoloma mougeotii | 2 | 1 | x | x | Agaricales | Agaricomycetes | SH213985.07FU |
| Entoloma neglectum | | 1 | | | Agaricales | Agaricomycetes | SH185816.07FU |
| Entoloma nitidum | | 9 | | | Agaricales | Agaricomycetes | SH187111.07FU |
| Entoloma prunuloides | 2 | 2 | x | x | Agaricales | Agaricomycetes | SH187841.07FU |
| Entoloma queletii | 1 | | x | | Agaricales | Agaricomycetes | |
| Entoloma rhombisporum | 2 | | x | x | Agaricales | Agaricomycetes | |
| Entoloma sodale | 1 | | x | | Agaricales | Agaricomycetes | |
| Entoloma turci | | 1 | | | Agaricales | Agaricomycetes | SH221240.07FU |
| Entoloma xanthochroum | 1 | | x | | Agaricales | Agaricomycetes | |
| Exidia repanda | 1 | | | | Auriculariales | Agaricomycetes | |
| Flammulaster limulatus | 1 | | | | Agaricales | Agaricomycetes | |
| Ganoderma adspersum | | 2 | | | Polyporales | Agaricomycetes | SH187221.07FU |
| Ganoderma pfeifferi | 1 | 2 | | x | Polyporales | Agaricomycetes | SH187235.07FU |
| Geastrum corollinum | | 1 | | | Geastrales | Agaricomycetes | SH211464.07FU |
| Gloeocystidiellum clavuligerum | 1 | | | | Russulales | Agaricomycetes | |

| | | | | | | | |
|---|---|---|---|---|---|---|---|
| Gloeoporus pannocinctus | 2 | | | x | Polyporales | Agaricomycetes | |
| Gloioxanthomyces vitellinus | 4 | 2 | x | x | Agaricales | Agaricomycetes | SH178713.07FU |
| Glutinoglossum glutinosum | 7 | | x | x | Geoglossales | Geoglossomycetes | |
| Gymnopus inodorus | 1 | | x | | Agaricales | Agaricomycetes | |
| Gyrodon lividus | 1 | | x | x | Boletales | Agaricomycetes | |
| Hebeloma fusisporum | 2 | | x | | Agaricales | Agaricomycetes | |
| Hebeloma vaccinum | 1 | | x | x | Agaricales | Agaricomycetes | |
| Helvella bicolor | | 1 | | | Pezizales | Pezizomycetes | SH174882.07FU |
| Hodophilus foetens | 1 | 1 | x | x | Agaricales | Agaricomycetes | SH523020.07FU |
| Hodophilus hymenocephalus | 1 | | x | x | Agaricales | Agaricomycetes | |
| Hohenbuehelia atrocoerulea | 1 | | | | Agaricales | Agaricomycetes | |
| Hohenbuehelia mastrucata | 2 | | | x | Agaricales | Agaricomycetes | |
| Hohenbuehelia petaloides | 1 | | x | x | Agaricales | Agaricomycetes | |
| Hygrocybe cantharellus | 13 | 8 | x | x | Agaricales | Agaricomycetes | SH221288.07FU |
| Hygrocybe coccineocrenata | 2 | | x | | Agaricales | Agaricomycetes | |
| Hygrocybe helobia | 3 | | x | x | Agaricales | Agaricomycetes | |
| Hygrocybe mucronella | 2 | | x | x | Agaricales | Agaricomycetes | |
| Hygrocybe phaeococcinea | 1 | | x | x | Agaricales | Agaricomycetes | |
| Hygrocybe punicea | | 3 | | | Agaricales | Agaricomycetes | SH199257.07FU |
| Hygrocybe quieta | | 1 | | | Agaricales | Agaricomycetes | SH485060.07FU |
| Hygrocybe reidii | 6 | 5 | x | x | Agaricales | Agaricomycetes | SH191907.07FU |
| Hygrocybe spadicea | 1 | | x | x | Agaricales | Agaricomycetes | |
| Hygrocybe splendidissima | 1 | 2 | x | x | Agaricales | Agaricomycetes | SH107463.07FU |
| Hygrocybe substrangulata | 1 | 1 | x | x | Agaricales | Agaricomycetes | SH208313.07FU |
| Hygrocybe turunda | | 7 | | | Agaricales | Agaricomycetes | SH221291.07FU |
| Hygrophorus mesotephrus | | 1 | | | Agaricales | Agaricomycetes | SH000341.07FU |
| Hygrophorus persoonii | | 1 | | | Agaricales | Agaricomycetes | SH202873.07FU |
| Hygrophorus piceae | 1 | | x | x | Agaricales | Agaricomycetes | |
| Hygrophorus unicolor | 1 | | x | | Agaricales | Agaricomycetes | |
| Hymenochaetopsis corrugata | 2 | | | x | Hymenochaetales | Agaricomycetes | |
| Hyphoderma macedonicum | 2 | | | | Polyporales | Agaricomycetes | |
| Hyphoderma medioburiense | 1 | | | x | Polyporales | Agaricomycetes | |
| Inocybe calamistrata | 1 | 2 | x | x | Agaricales | Agaricomycetes | SH188895.07FU |
| Inocybe flavella | | 2 | | | Agaricales | Agaricomycetes | SH032895.07FU |
| Inocybe hystrix | | 1 | | | Agaricales | Agaricomycetes | SH218251.07FU |
| Inocybe margaritispora | 1 | | x | x | Agaricales | Agaricomycetes | |
| Inocybe pelargonium | 1 | | x | x | Agaricales | Agaricomycetes | |
| Inocybe praetervisa | 2 | | x | x | Agaricales | Agaricomycetes | |
| Inonotus hispidus | | 3 | | | Hymenochaetales | Agaricomycetes | SH188978.07FU (as Agaricomycetes sp) |
| Ischnoderma resinosum | 2 | | | x | Polyporales | Agaricomycetes | |
| Jaapia ochroleuca | 11 | | | | Boletales | Agaricomycetes | |
| Kneiffiella alutacea | 6 | | | | Hymenochaetales | Agaricomycetes | |

| | | | | | | | |
|---|---|---|---|---|---|---|---|
| Kneiffiella barba-jovis | 7 | | | x | Hymenochaetales | Agaricomycetes | |
| Lactarius azonites | | 4 | | | Russulales | Agaricomycetes | SH221822.07FU (as Lactarius acris) |
| Lactarius mammosus | | 8 | | | Russulales | Agaricomycetes | SH176401.07FU |
| Lactarius trivialis | 1 | | x | x | Russulales | Agaricomycetes | |
| Lactarius uvidus | 1 | | x | x | Russulales | Agaricomycetes | |
| Lentaria byssiseda | 2 | | | x | Gomphales | Agaricomycetes | |
| Lentinellus flabelliformis | 1 | | | x | Russulales | Agaricomycetes | |
| Lentinellus ursinus | 1 | | | x | Russulales | Agaricomycetes | |
| Lepiota ochraceofulva | 1 | | x | | Agaricales | Agaricomycetes | |
| Lepiota pseudolilacea | 2 | 1 | x | x | Agaricales | Agaricomycetes | SH180338.07FU |
| Lepiota tomentella | 1 | | x | x | Agaricales | Agaricomycetes | |
| Lindtneria trachyspora | 4 | | x | x | Agaricales | Agaricomycetes | |
| Lyophyllum leucophaeatum | | 1 | | | Agaricales | Agaricomycetes | SH533909.07FU |
| Microglossum olivaceum | 1 | | x | x | Leotiales | Leotiomycetes | |
| Microglossum viride | 2 | 2 | x | x | Helotiales | Leotiomycetes | SH212648.07FU |
| Mitrula paludosa | | 2 | | | Helotiales | Leotiomycetes | SH211648.07FU |
| Mycena clavata | 3 | | | | Agaricales | Agaricomycetes | |
| Mycena concolor | 2 | | x | | Agaricales | Agaricomycetes | |
| Mycena meliigena | 2 | | | | Agaricales | Agaricomycetes | |
| Mycena picta | 2 | | x | | Agaricales | Agaricomycetes | |
| Nemania carbonacea | 2 | | | | Xylariales | Sordariomycetes | |
| Nemania diffusa | 4 | | | x | Xylariales | Sordariomycetes | |
| Neofavolus suavissimus | 2 | | | x | Polyporales | Agaricomycetes | |
| Neohygrocybe ingrata | | 2 | | | Agaricales | Agaricomycetes | SH193313.07FU |
| Neohygrocybe nitrata | | 1 | | | Agaricales | Agaricomycetes | SH495419.07FU |
| Peniophorella guttulifera | 3 | | | x | Hymenochaetales | Agaricomycetes | |
| Phlebia subochracea | 1 | | | x | Polyporales | Agaricomycetes | |
| Pholiota henningsii | 1 | | x | x | Agaricales | Agaricomycetes | |
| Pholiota tuberculosa | 4 | | | x | Agaricales | Agaricomycetes | |
| Plicatura crispa | 4 | | | x | Agaricales | Agaricomycetes | |
| Pluteus exiguus | 2 | | | x | Agaricales | Agaricomycetes | |
| Pluteus hispidulus | 1 | | | | Agaricales | Agaricomycetes | |
| Pluteus leoninus | 2 | | | x | Agaricales | Agaricomycetes | |
| Pluteus pellitus | | 1 | | | Agaricales | Agaricomycetes | SH185148.07FU |
| Poronia punctata | 1 | | x | x | Xylariales | Sordariomycetes | |
| Porotheleum fimbriatum | 1 | | | x | Agaricales | Agaricomycetes | |
| Psathyrella suavissima | | 2 | | | Agaricales | Agaricomycetes | SH208812.07FU |
| Pseudotricholoma metapodium | 1 | | x | x | Agaricales | Agaricomycetes | |
| Psilocybe turficola | 1 | | x | x | Agaricales | Agaricomycetes | |
| Ramariopsis pulchella | 4 | | x | | Agaricales | Agaricomycetes | |
| Rhodophana nitellina | 1 | | x | x | Agaricales | Agaricomycetes | |
| Russula albonigra | | 1 | | | Russulales | Agaricomycetes | SH219262.07FU |

| | | | | | | | |
|---|---|---|---|---|---|---|---|
| Russula alnetorum | 2 | 4 | x | x | Russulales | Agaricomycetes | SH190470.07FU |
| Russula anthracina | 1 | | x | | Russulales | Agaricomycetes | |
| Russula pelargonia | 3 | | x | x | Russulales | Agaricomycetes | |
| Russula puellula | | 5 | | | Russulales | Agaricomycetes | SH219862.07FU (as Russula puellaris) |
| Russula sanguinea | 1 | | x | x | Russulales | Agaricomycetes | |
| Simocybe sumptuosa | 2 | | | | Agaricales | Agaricomycetes | |
| Steccherinum subcrinale | 1 | | x | | Polyporales | Agaricomycetes | |
| Stypella dubia | 3 | | | x | Auriculariales | Agaricomycetes | |
| Stypella subgelatinosa | 1 | | | | Auriculariales | Agaricomycetes | |
| Tomentella lateritia | 2 | 9 | x | x | Thelephorales | Agaricomycetes | SH185279.07FU |
| Tomentella pilosa | | 2 | | | Thelephorales | Agaricomycetes | SH177790.07FU |
| Tomentella umbrinospora | 1 | 1 | x | x | Thelephorales | Agaricomycetes | SH185243.07FU |
| Trechispora silvae-ryae | 2 | | x | | Trechisporales | Agaricomycetes | |
| Tremellodendropsis tuberosa | 1 | | x | x | Tremellodendropsidales | Agaricomycetes | |
| Tricholoma inamoenum | 1 | 1 | x | x | Agaricales | Agaricomycetes | SH190400.07FU |
| Tricholoma umbonatum | | 1 | | | Agaricales | Agaricomycetes | SH493974.07FU |
| Xenasma pruinosum | 1 | | | x | Polyporales | Agaricomycetes | |
| Xenasma pulverulentum | 4 | | | | Polyporales | Agaricomycetes | |
| Xerula pudens | 1 | | x | x | Agaricales | Agaricomycetes | |
